# Cryo-EM reveals the mechanism of DNA compaction by *Mycobacterium smegmatis* Dps2

**DOI:** 10.1101/2023.01.16.523357

**Authors:** Priyanka Garg, Thejas Satheesh, Mahipal Ganji, Somnath Dutta

## Abstract

<u>D</u>NA-binding protein under starvation (Dps), is a miniature ferritin complex which plays a vital role in protecting bacterial DNA during starvation for maintaining the integrity of bacteria from hostile conditions. *Mycobacterium smegmatis* is one such bacteria that express MsDps2, which binds DNA to protect it under oxidative and nutritional stress conditions. Several approaches, including cryo-electron tomography (Cryo-ET), were implemented to identify the structure of the Dps protein that is bound to DNA. However, none of the structures of the Dps-DNA complex was resolved to high resolution to be able to identify the DNA binding residues. In this study, we implemented various biochemical and biophysical studies to characterize the DNA protein interactions of Dps protein. We employed single-particle cryo-EM-based structural analysis of MsDps2-DNA and identify that the region close to N-terminal confers the DNA binding property. Based on cryo-EM data, we performed mutations of several arginine residues proximal to DNA binding region, which dramatically reduced the MsDps2-DNA interaction. In addition, we demonstrated the proposed model for DNA compaction during lattice formation. We also pinpointed arginine residues, which are responsible for DNA binding in lattice arrangement of MsDps2. We performed single-molecule imaging experiments of MsDps2-DNA interactions that corroborate well with our structural studies.

## Introduction

DNA-protein interaction in a cellular organism plays a significant spatial and temporal role concerning the organism’s development (von Hippel & McGhee, 1972). Proteins bind DNA in either sequence-specific or non-specific manner (Strauch, 2001). The sequence specificity might involve the recognition of a defined sequence or a consensus sequence. Particular structural motifs on a given protein usually make a direct contact with DNA in the case of sequence-specific binding (Ferraz *et al*., 2021, Mangiarotti *et al*., 2009). While some proteins, such as RNA polymerase or DNA polymerase, have a single DNA binding site, others, such as transcription regulatory factors, involve multiple repetitive binding motifs for recognizing the specific DNA sequence. Some structural motifs involved in DNA binding are the helix-turn-helix motif, zinc-coordinating motif, leucine zipper motif, and other alpha helix and the β-sheet motifs (Luscombe *et al*., 2000, Struhl, 1989).

A non-specific DNA-protein interaction, on the other hand, predominantly involves ionic interactions with the sugar-phosphate backbone of DNA (Luscombe *et al*., 2000). Most proteins that show site-specific binding with DNA also display significant non-specific DNA affinity. DNA-binding proteins with multiple binding domains leads to large supramolecular structures (Ganguly *et al*., 2012). Such multiple binding interactions occur primarily in the case of genome organization or physical protection of DNA from environmental factors. For example, Nucleoid-Associated Proteins (NAPs) in bacteria help in chromatin structure, and DNA replication, repair, protection and transcription (Hołówka & Zakrzewska-Czerwińska, 2020). Each of the NAPs distinctly binds DNA to serve their function. Nucleoid-associated proteins (NAPs), in bacteria display non-specific interactions which tightly compact DNA (Hołówka & Zakrzewska-Czerwińska, 2020). This binding may lead to remodeling of the nucleoid’s structure which might in turn alter the gene expression. Therefore, it is crucial to investigate the nature of complexes formed between proteins and DNA. Over the past decade, we have witnessed a significant expansion in the determination of high-resolution structures of DNA-binding proteins (Luscombe *et al*., 2000). The structural characterization of protein-DNA complexes have provided considerable insight into the stereochemical principles of binding.

DNA-binding protein from stationary phase cells (Dps) is one such protein belonging to the family of NAPs and has been shown to play an essential role in protecting microorganisms from oxidative and nutritional stress (Fenton, 1894). It also helps in Iron oxidation and storage (Martinez & Kolter, 1997). Dps is known to interact with DNA in a sequence-independent manner. The structures of Dps proteins reveal a subunit core consisting of a four-helix bundle and a long-loop containing a small helix in the middle, identical to that in Dps from other sources (Haikarainen & Papageorgiou, 2010). The surface charge of Dps is predominantly negatively charged. The DNA binding ability varies in all members of this family since the establishment of effective electrostatic interactions with the negatively charged DNA backbone depends on the presence of specific structural elements on the Dps molecule (Stillman *et al*., 2005, Hitchings *et al*., 2014). There is a difference in the DNA binding ability among the homologs of Dps depending on the variability of the number of positive charges present in N- and C-terminal tails. Three DNA binding signatures are based on the presence and length of N- and C-terminal loops. For instance, a long N-terminal tail in *E. coli* Dps has three lysine residues that can interact with the phosphate groups on DNA (Grant *et al*., 1998). A C-terminal tail executes the DNA binding function in Mycobacterium Smegmatis Dps1 (MsDps1) (Roy *et al*., 2007). Unlike the *E. coli* one, MsDps1 also has a short N-terminal tail that is imperative for multimeric assembly. The third DNA binding signature is the surface of the Dps molecule like *H. pylori* Dps, also named HPNAP as this Dps has no Nor C-terminal tail.(Ceci *et al*., 2007).

Scrutinizing of these structural elements of Dps molecules used to bind DNA provides a complete picture of the modes of interaction between the two macromolecules. Thus, with the presence or absence of these signatures in the sequence of a given Dps protein can be used to predict its DNA binding ability with reasonable confidence. Indeed, Dps proteins with a shorter length of N-terminus, like *L. innocua* Dps, *B. anthracis* Dlp-1 and Dlp-2, and *Campylobacter jejuni* (*Cj-DPS*), do not interact with DNA, according to predictions. The flexibility of the N-, and C-terminus tails are also vital as the immobilization of the N-terminal tail of *A. tumefaciens* Dps results in abolishing its DNA binding activity (Ceci *et al*., 2003). The role played by the multiplicity of positively charged residues and by the N- or C-terminus mobility in determining the strength and pH dependence of the interaction was evaluated in later studies (Dubrovin *et al*., 2021, Kamyshinsky *et al*., 2019). Various techniques, namely electrophoretic mobility shift assays and atomic force microscopy, negative staining TEM and cryo-EM tomography, were used to probe the interaction of Dps–DNA (Tereshkin *et al*., 2019, Dadinova *et al*., 2021, Dubrovin *et al*., 2021, Kamyshinsky *et al*., 2019).

The non-pathogenic mycobacterial species *M. smegmatis* has two Dps homologs, MsDps1 and MsDps2 (Gupta & Chatterji, 2003). These are differently regulated in the cells, where MsDps1 is expressed in the stationary phase and is controlled by the extracellular sigma factors sigF and sigH, whereas MsDps2 is constitutively expressed by RNA polymerase under the control of sigma factors sigA and sigB (Chowdhury *et al*., 2007). A variation is the *M. smegmatis* Dps protein MsDps1, which seemed to bind DNA through a mobile and positively charged C-terminus with three lysines and two arginines. Deleting 16 C-terminal residues resulted in a loss of DNA binding activity (Roy *et al*., 2007). Deleting the entire C-terminal tail resulted in an open decamer assembly, indicating that the C-terminal tail has a role in DNA binding and oligomerization. Unlike MsDps1, MsDps2 does not have a long C-terminal tail. It has an N-terminal tail deletion of which does not entirely abolish the DNA activity.

MsDps2 forms a stable dodecamer which possesses the ability to carry out ferroxidation activity (Roy *et al*., 2008). When a microorganism is under stress, it releases reactive oxidative species, which harm cellular metabolism. The dodecameric form can thus provide physical protection to DNA from oxidative stress and help the organism’s survival. Although the crystal structure of the MsDps2 dodecamer has been solved at 2.4 Å resolution, no atomic-scale structure of MsDps2–DNA assemblies currently exist, and little is known about complex formation and DNA protection mechanism (Roy *et al*., 2008, Saraswathi *et al*., 2009).

In the present study, we have used biophysical and microscopy techniques to probe the DNA binding property of the MsDps2 and provide insights into understanding DNA protection mechanism. We used gel retardation assay in agarose gels to assess the binding of MsDps2 with different types of DNA. In this assay, MsDps2 forms a large multiprotein complex with DNA and shows a few intermediate-sized complexes. Further, we have probed the DNA binding activity by using negative staining TEM. To decipher the structure of MsDps2-DNA complex, we performed cryo-EM. We gathered insights into the binding site as well as the residues required for the DNA binding property of MsDps2. After this, we generated different truncates of MsDps2 and simultaneously mutated important residues. By performing EMSA and single-molecule experiments, we delineate the effect of these critical DNA binding signatures and residues. In addition, we demonstrated a model for DNA compaction during lattice formation.

## Material and Methods

### Cloning of different constructs of MsDps2

MsDps2 clone was gifted to us by Prof. Dipankar Chatterji, and the cloning protocol was described in Roy *et al*., 2008, Journal of Molecular Biology. Briefly, the gene for MsDps2 was cloned from the *M. smegmatis* genomic DNA into pET21b using enzymes Nde I (NEB #R0111S) and HindIII-HF (NEB #R3104S). MsDps2 plasmid was used as a template, and several other constructs were generated for this study: one is with the deletion of 15 amino acids from the N-terminal of MsDps2 (ΔN_15_MsDps2).Additionally, we generated a variant with multiple point mutations (Arg4Glu, Arg5Glu, Arg102Glu, and Arg114Glu) using similar approaches. The primers used for generating the truncates and mutants are listed in Table S1.

### Expression and purification of recombinant Msdps2 and its constructs

*E. coli* BL21(DE3) cells harbouring the pET21b-MsDps2 plasmid and its different constructs were grown in 1.6 litre of liquid Luria Bertani broth (HiMedia Laboratories, Mumbai, India) containing ampicillin (100 μg/ml) at 37 °C to an optical density of 0.6 at 600 nm (OD_600_). After adding 1 mM isopropyl-β-D-thiogalactopyranoside (IPTG) to induce the expression of the MsDps2 and its variants gene, the culture was incubated further for 3–4 h. After that, the cells were harvested (50,000 rpm for 20 min), suspended in 40 ml of lysis buffer 50 mM Tris–HCl (pH 7.9), 1 M NaCl and disrupted by sonication. The lysate was centrifuged at 12,000 rpm for 45 min. Then, the proteins were separated by precipitating the nucleic acids using 5% PolyEthyleneImine (PEI). Five percent PEI stock solution was added progressively into the extract by gently stirring until the final concentration of PEI was 0.6%. During this process, the nucleic acids were precipitated. The precipitation was removed by centrifugation at 17,300*g* for 10 min, leaving the proteins in the supernatant.

The supernatant was precipitated using 30% ammonium sulphate cuts. At the 30% cut, MsDps2 protein was precipitated and resuspended in the buffer. This solubilized protein was dialyzed overnight against 20 mM Tris–HCl (pH 7.0) and 50 mM NaCl and then loaded onto an anionic diethylaminomethyl –Sephacel (Sigma-Aldrich I6505) equilibrated with the same buffer. MsDps2 was eluted with 200 mM NaCl. The eluted proteins were injected and purified on SEC (size exclusion chromatography) column Superdex 200 10/300 GL (GE Healthcare Life Sciences, Piscataway, NJ, USA) against a buffer containing 20 mM Tris-HCl (pH 7.5) and 200 mM NaCl. Different fractions collected during protein purification were analyzed by 10% SDS-PAGE.

### Gel retardation assay

The Electrophoretic Mobility Shift Assay (EMSA) was performed by using linear (600bp, 200bp) and plasmid (3kbp) in 20 mM Tris–HCl (pH 7.5), 75 mM NaCl. 50 ng of each DNA was incubated with varying the concentration of MsDps2 (1 μM −12 μM) and its constructs at room temperature for 5 min in 20μl of total reaction volume. The complex was then resolved on a 0.6% agarose gel in 1× TAE [Tris-acetate–ethylenediaminetetraacetic acid (EDTA)] buffer consisting of 40 mM Tris-acetate, 20 mM sodium acetate and 1 mM EDTA (pH 8.0). The electrophoresis was carried out at a constant voltage of 50 V. The gel was stained with ethidium bromide and observed under UV light.

Similarly, the reaction with fluorescent 50bp of MsDps2 at different concentrations was loaded on Native PAGE gel, stained with ethidium bromide, and observed under UV light.

### Binding affinity studies by Microscale Thermophoresis (MST)

MST was performed for binding affinity measurement of MsDps2 with DNA, where the protein was labelled with Monolith Protein labelling kit RED-NHS (NanoTemper Technologies, Munich, Germany). In the binding assay, the protein concentration for native was kept constant. The proteins were incubated with 16 twofold serial dilutions of the ligand DNA. Samples were loaded on MST Premium Coated Monolith™ NT.115 capillaries, and measurements were done using Monolith NT.115pico. Further data analysis was done using Monolith Affinity Analysis software (version 2.3).

### Negative stain single-particle Electron Microscopy (EM)

Purified MsDps2 and its variants were incubated with various types of DNA and visualized by negative staining TEM to observe the interaction of protein and DNA. All the samples were prepared by conventional negative staining methods (Kar *et al*., 2022). A carbon-coated copper grid (EM grid, 300 mesh; TedPella) was glow-discharged (GloQube glow discharge system, Quorum) for 30 s at 20 mA. A total of 3.5 μl sample (0.01 mg/ml) was added to the grid for 30 s. The extra sample was blotted out. Negative staining was performed using 1% uranyl acetate (98% uranyl acetate; ACS Reagent, Polysciences Inc. Warrington, PA, USA) solution for 20 s. The grid was air-dried. The negatively stained sample for native MsDps*2* and MsDps2 with 100bp DNA were visualized at room temperature using Tecnai T12 electron microscope equipped with a LaB_6_ filament operated at 120 kV, and images were recorded using a side-mounted Olympus VELITA (2000 × 2000) charge-coupled device camera at a magnification of ×220,000 (2.54 Å per pixel). MsDps2 with plasmid DNA and MsDps2 with 600bp DNA were visualized at room temperature using a Talos L120C transmission electron microscope (Thermo Fisher Scientific) equipped with Ceta (4000 × 4000) camera. Images were recorded at a magnification of ×92,000 (1.52 Å per pixel). Micrographs were evaluated with EMAN 2.1 (Tang *et al*., 2007). Protein particles were manually picked and extracted with a box size of 160 Å from the raw micrographs using e2boxer.py (Ludtke, 2012). The two-dimensional reference-free classification of the extracted particles was performed using e2projectmanager.py (EMAN2.1 software). Similarly, it was done for all other MsDps2-DNA complexes and other MsDps2 constructs.

### Cryo-EM Grid preparation

R 1.2/1.3 (Quantifoil) (Electron Microscopy Sciences) 300-mesh copper grids were glow-discharged for 90 s at 20 mA using Quorum GlowQube before sample preparation.3 μl of 0.1 mg/ml purified MsDps2 protein, MsDps2-DNA complex and ΔN_15_MsDps2-DNA (50 mM Tris buffer pH 7.9 and 100 mM NaCl) were applied to glow discharged grids. Grids were further incubated for 10 s at > 100% humidity and blotted for 8 seconds before freezing. The grids were plunge-frozen in liquid ethane using Thermo Fisher Scientific Vitrobot Mark IV.

### Cryo-EM data collection of Wild Type MsDps2 and its different constructs

For high-resolution cryo-EM structural characterization, the MsDps2 dataset was collected on Thermo Scientific^™^ Talos Arctica Transmission Electron Microscope at 200 kV at a nominal magnification of 72,000 x, equipped with a K2 Summit Direct Electron Detector (Gatan Inc). Latitude-S automatic data collection software (Gatan Inc) was utilized to collect the 404 movies at a pixel size of 0.74 Å with a total electron dose of about 40 e-/Å^2^ at the defocus range of −0.75 and −2.25 μm, under a calibrated dose of about 2 e-/Å^2^ per frame. Movies were recorded for 8 secs with 20 frames. Similar approaches were implemented to collect data for MsDps2 with 600bp DNA and ΔN15MsDps2-DNA with 50bp DNA. However, 1091 movies of MsDps2 with 600bp DNA was acquired at a nominal magnification of 42,200x and a pixel size of 1.17 Å, whereas 838 movies of ΔN15MsDps2-DNA with 50bp DNA was collected at a nominal magnification of 54,000 x (pixel size of 0.92 Å) (Kumar *et al*., 2021). Moreover, electron dose, number of frames, and defocus values were same for all three samples.

### Cryo-EM data processing of different constructs of MsDps2

Data processing was primarily performed using cryoSPARC 3.3.1 (Punjani *et al*., 2017a). Initially, beam-induced motion correction of the individual movies was performed using Patch motion correction (Grant *et al*., 2018). All the motion-corrected micrographs were manually curated using cisTEM software package (Grant *et al*., 2018), and the best micrographs were considered for further processing. Contrast transfer function (CTF) was estimated using patch CTF (Zhang, 2016).

For MsDps2 with 600bp DNA, 973,755 particles were automatically selected using the cryoSPARC template picker. The particles were extracted with a box size of 256 pixels. Three rounds of reference-free 2D classification were carried out with 136 Å particle diameter to remove bad particles. 417,449 particles were selected, and the ab initio initial model was generated using 12,132 particles with C1 symmetry. Further, these particles were separated into 10 classes by performing heterogeneous refinement by imposing no symmetry with particle diameter 136 Å. Of the 10 classes, class 1, class 4 and class 6 showed better secondary structure features than other classes. These classes were selected for uniform homogeneous refinement. The map was further refined by global CTF refinement. The resolution of the 3D map was estimated at FSC 0.143. Later, the focused refinement of the core was performed by imposing no symmetry with a particle diameter of 120 Å, and particles were classified into three classes. Class 1 was further refined by uniform homogeneous refinement by imposing T symmetry. The map was further refined by global CTF refinement. The resolution of the 3D map was estimated at FSC 0.143. The heterogeneous refinement was again performed by keeping the same particles and other parameters except for changing the particle diameter to 200 Å. Class 2 was taken further for refinement, as mentioned above (Supplementary figure 3).

For wild-type MsDps2, 289,129 particles were automatically selected using the cryoSPARC template picker. The particles were extracted with a box size of 328 pixels. Three rounds of reference-free 2D classification were carried out with 120 Å particle diameter to remove bad particles. 219,501 particles were selected, and ab initio initial model was generated with T symmetry with 11,289 particles. The particles were classified into 3 classes by performing heterogeneous refinement. Class 3 was taken further for uniform refinement with T symmetry. The map was further refined by global CTF refinement (Punjani *et al*., 2017b) and also for Ewald sphere correction (Zivanov *et al*., 2018). The resolution of the 3D map was estimated at FSC 0.143 (Supplementary figure 4).

For ΔN_15_MsDps2-DNA with 50bp DNA, 177,500 particles were automatically selected using the cryoSPARC template picker. The particles were extracted with a box size of 320 pixels. Three rounds of reference-free 2D classification were carried out with a 160 Å particle diameter to remove bad particles. 41,049 particles were selected, and an ab initio initial model was generated without symmetry. Further, these particles were sorted into 3 classes by performing heterogeneous refinement by imposing no symmetry of the three ab initio models. Class 3 were selected for uniform homogeneous refinement. Later, the focused refinement of the core was performed by imposing no symmetry with a particle diameter of 120 Å, and particles were classified into three classes. Class 1 was further refined by uniform homogeneous refinement by imposing T symmetry. The map was further refined by global CTF refinement. The resolution of the 3D map was estimated at FSC 0.143 (Supplementary figure 7).

### Phenix refinement and Coot

For the model building of MsDps2, the available crystal structure of MsDps2 (PDB-2Z90) was used as a template (Roy *et al*., 2008). The biological assembly model of the MsDps2 was manually fit into the cryo-EM map of MsDps2 using UCSF Chimera and Chimera X (Pettersen *et al*., 2004, Goddard *et al*., 2018). The model of MsDps2 was further inspected manually using Coot (Emsley *et al*., 2010). This model was refined with Phenix using real-space refinement (Afonine *et al*., 2018). For docking of the MsDps2-DNA complex, the DNA structure was generated by 3D DNA (Pettersen *et al*., 2004). The DNA and the atomic model of MsDps2 were manually fit into the cryo-EM map of MsDps2-DNA.

The model of ΔN_15_MsDps2 was generated by deleting 15 residues from each monomer of the atomic model of MsDps2.This model was used as a template and fit into the cryo-EM density of ΔN_15_MsDps2. This model was further refined with Phenix using real-space refinement (Afonine *et al*., 2018). The DNA and atomic model of ΔN_15_MsDps2 were docked into the cryo-EM density of ΔN_15_MsDps2-DNA. Figures were made with Chimera and Pymol (DeLano, 2002, Pettersen *et al*., 2004).

### Slide preparation for single-molecule fluorescence experiments

Glass slides with microfluidic channels were prepared in-house. For this, we drilled holes on each side of the glass slide (VWR #631-1550) using a Dremel drill (#Dremel 4000) with a diamond head drill bit of 2 mm diameter for the preparation of microfluidic channels. The slides and the coverslips (VWR #631-0147) were initially cleaned in a series of steps by rinsing and sonicating using MilliQ water, acetone (SRL #31566), and potassium hydroxide KOH (BDH #296228). Acid piranha solution (3:1 of H_2_SO_4_:H_2_O_2_) was used for further cleaning, and the etching additionally generates hydroxyl groups on the surface. It is essential for functionalizing the slides and coverslips with an amine group. An aminosilanization reaction mixture is prepared using a 2:1:20 ratio of APTES (3-Aminopropyl triethoxysilane, SRL #33993), glacial acetic acid (SDFCL #20001), and methanol, respectively. Slides and coverslips were then incubated for around 25 minutes in the freshly prepared aminosilanization mixture. After aminosilanisation, the surfaces were peg-passivated using a mixture of 1:20 ratio of biotin-PEG-SVA (Succinimidyl Valerate) (Laysan Bio #Biotin-PEG-SVA-5000) to mPEG-SVA (Laysan Bio #mPEG-SVA-5000) dissolved at 20 mM concentration in a freshly prepared 0.1 M of sodium bicarbonate buffer (Sigma #S5761) of pH 8.5. A 60 μL of this mixture was then sandwiched between a glass slide and a cover slip. These sandwiches were incubated in a dark and humid environment overnight. The slides and coverslip were disassembled, rinsed with MilliQ water, and dried using compressed nitrogen gas. An additional round of PEGylation was carried out for further passivation of the surface before using for single-molecule imaging experiments. For this purpose, we used MS(PEG)_4_ (Thermo Scientific #22341). A 6 μL of 250 mM MS(PEG)_4_ was dissolved in 54 μL of the sodium bicarbonate buffer.

### Preparation of biotin-labelled DNA

The cohesive ends of lambda phage DNA (SRL #67291) were filled using a Klenow fragment (NEB #M0210L) with dATP, dGTP, dCTP and biotin-11-dUTP (Jena Bioscience #NU-803-BIOX-S) to generate DNA with biotin labelling on both ends. The labelled DNA was then purified using a PCR clean up kit (Qiagen #28104) to remove the free nucleotides. The biotin-modified DNA was digested using a KasI restriction enzyme (NEB #R0544S) that produced two fragments of 24 kb and 24.5 kb, respectively. This preparation resulted in similar-sized DNA molecules having a single biotin moiety at one end.

### Flow cell preparation

The flow cell used in the experiment was prepared by sandwiching a double-sided tape between the PEG-passivated glass slide and coverslip. The ends of the channels were sealed with epoxy glue. The drilled holes in the glass slide enable buffer exchange. These channels were first incubated with 20 μL of 4 % (w/v) BSA (Sigma-Aldrich #A2153) for 5 minutes. The introduction of BSA was done using a pipette tip plugged into one of the holes of the channel, acting as an inlet. The other hole attached to a syringe through polypropylene tubing served as an outlet. The channels were washed with 500 μL of T50 buffer (50 mM Tris-HCl pH 7.5, 50 mM NaCl, and 0.2 mM EDTA). Next, a 20 μL of 0.2 % (v/v) Tween20 was flowed into the channel and incubated for 5 minutes, followed by a 500 μL T50 wash. We incubated with 0.5 mg/mL streptavidin (SRL #87610) in T50 buffer for 10 minutes. Finally, we washed the flow cell with 500 μL of T50 buffer before immobilising biotin-labelled DNA.

### Preparation of oxygen scavenging system

PCD (Protocatechuate 3,4-dioxygenase; Sigma #P8279-25UN) was dissolved in a buffer containing 100 mM Tris–HCl, pH 8.0, 1 mM EDTA, 50 mM KCl and 50% glycerol (Sigma #G5516-1L) to make 20X PCD with a final concentration of 6 μM. A 154 mg of PCA (Protocatechuic acid/ 3,4-Dihydroxybenzoic acid; Sigma #37580-100G-F) was dissolved in 10 ml of HPLC grade water adjusted to pH 9.0 using 1 M NaOH (SRL #96311) to make 40X PCA. Trolox (6-hydroxy-2,5,7,8-tetramethylchroman-2-carboxylic acid; Sigma #238813-1G) was dissolved in 3.2 mL of HPLC grade water, 345 μL of 1M NaOH, and 430 μL of methanol to make a 100X Trolox.

### Single-molecule imaging

The imaging was done on a Nikon Ti2 eclipse microscope under total internal reflection mode. A 30 μL of the 1-5 pM of DNA was flowed in at a constant rate of 25 μL/min into the channel with the help of a motorized peristaltic pump. Once it was given sufficient time to bind to the streptavidin-coated glass surface, the unbound DNA was washed off with 50 μL of imaging buffer. The imaging buffer contained the T50 buffer along with 50 nM of Sytox Orange (Thermo Scientific #S11368), which was used to stain the DNA molecules, and an oxygen scavenging system (1X PCA, 1X PCD, 1X Trolox). This resulted in a distribution of single tethered DNA molecules on the surface, which could be imaged under a microscope. The sample was illuminated using a 561 nm wavelength laser which optimally excites Sytox Orange. The acquisition was done at 100 msec/frame for 30 sec to capture the single-tethered DNA molecules before introducing the protein. The appropriate concentration of MsDps2 protein of 50 μL has then flowed inside the channel. The DNA molecules were imaged continuously for one minute.

## Results

In this current study, we targeted to characterize the three-dimensional structure of MsDps2 with DNA to identify the DNA protection mechanism of bacteria during starvation. Additionally, we wanted to identify the critical amino acid residues responsible for MsDps2 for DNA binding. To achieve this, we first set to purify the MsDps2 protein from recombinant *E. coli* without any DNA contamination.

### Purification and biophysical characterization of DNA-free MsDps2

Previous studies showed that routine MsDps2 purification from *E. coli* BL21 cells resulted in bacterial DNA contamination (Saraswathi *et al*., 2009). Since we are specifically interested in deciphering the interactions between MsDps2 with DNA, removing the DNA in the early stage of MsDps2 purification is crucial. After expressing MsDps2 in *E. coli* BL21 cells, the cellular lysis was subjected to PolyEthyleneImine (PEI) precipitation before carrying Msdps2 protein purification, as PEI can remove the DNA contamination. Further, we analyzed the purified MsDps2 using analytical size exclusion chromatography (SEC) under native conditions to obtain information about the protein’s degree of oligomerization and homogeneity. The protein eluted at 68 ml as a dodecamer, corresponding to a molecular weight of 218 kDa (Figure 1A). We visualized the purified protein by SDS-PAGE and agarose gel. The SDS-PAGE showed a pure single band corresponding to a size of 18 kDa (Figure 1B).

**Figure 1:**
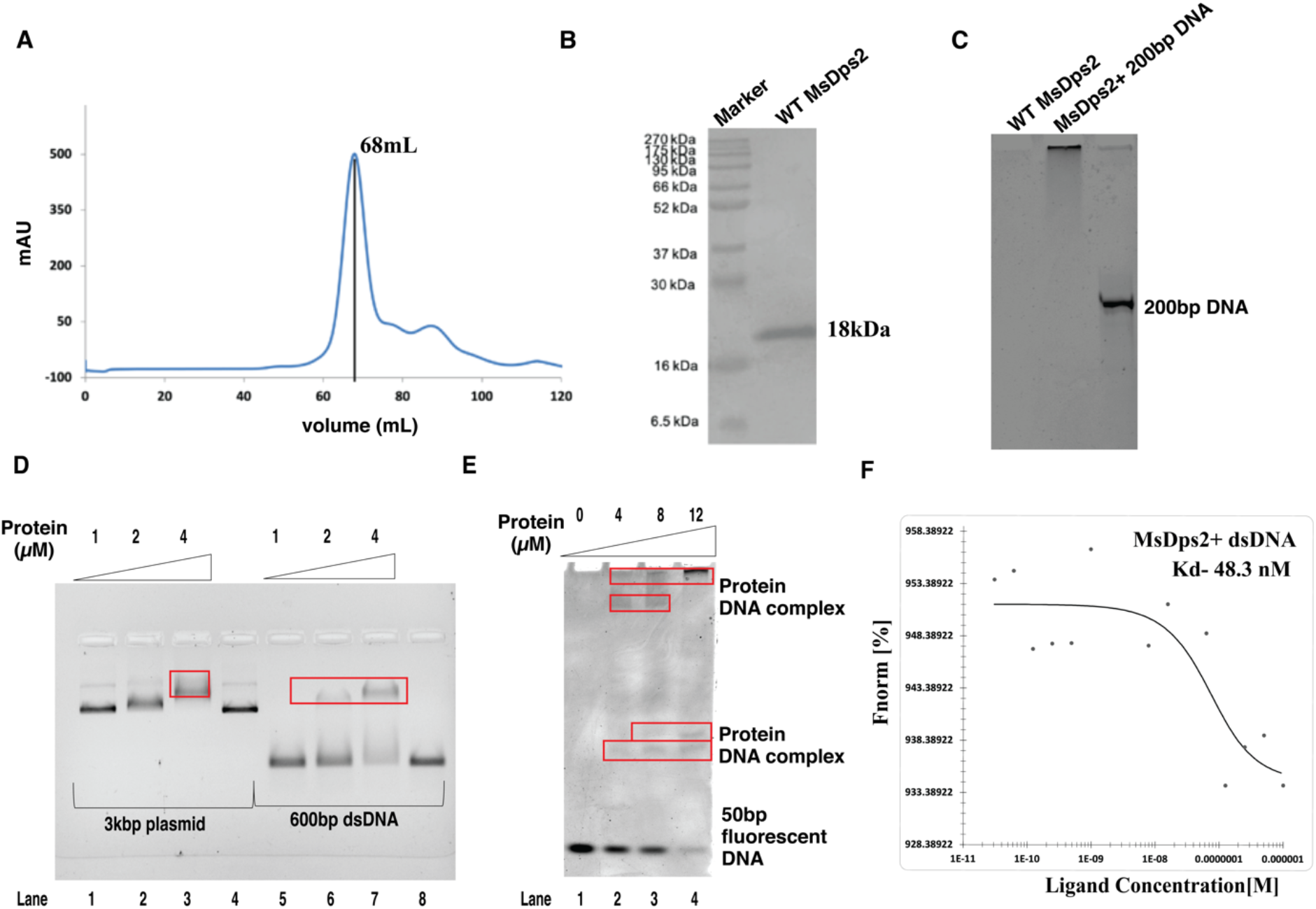
Biophysical characterisation and DNA binding activity of MsDps2: (A) SEC profile of Wild Type MsDps2 mAU, milli–Arbitrary Units (B) SDS PAGE analysis of MsDps2 where molecular weight of monomeric unit is 18kDa. (C) Agarose gel retardation assay: Lane 1 corresponds to only wild type MsDps2;Lane2 corresponds to 200bp DNA incubated with MsDps2; Lane3 corresponds to linear 200bp DNA. (D) Agarose gel retardation assay. Lane 1-3 corresponds to 3kbp plasmid DNA incubated with MsDps2 with increased concentration, Lane 4 is only 3kbp plasmid DNA whereas Lane 5-7 corresponds to 600bp linear DNA with MsDps2 with increased concentration, Lane 8 is only 600bp linear DNA. (E) DNA binding activity by using Native PAGE with fluorescent labelled 50bp DNA. (F) MST showing binding affinity of MsDps2 with DNA.

Further, no band of DNA was visible in agarose gel upon electrophoresion of the purified protein, suggesting that the protein was free of any DNA contamination (Figure 1C). The DNA-free MsDps2 enabled us to investigate MsDps2-DNA interactions.

### Biochemical characterization of DNA-Protein interactions

We then established the binding affinity of the purified MsDps2 to DNA. For this, we utilized Electrophoretic Mobility Shift Assay (EMSA) that detects DNA-protein interactions based on the retardation of the migration of complexes’ size. DNA mobility would be severely affected based on the number of proteins bound to DNA. Thus, we performed EMSA using MsDps2 with different types of DNA, such as double-stranded linear 200bp, 600 bp and 3 kb plasmid DNA. We incubated varied concentrations of MsDps2 with each DNA substrate and migrated on agarose gel under non-denaturing conditions. The results suggested that mobilities of DNA were severely affected in the presence of MsDps2 (Figure 1C-D), likely resulting from the strong affinity of MsDps2 to DNA. We note that there was no sequence similarity among these DNA substrates. Yet, the EMSA studies showed that the mobility of different types of DNA was significantly reduced in the presence of MsDps2. This indicates that MsDps2 binds DNA independent of its sequence, size, and topology. Thus, DNA sequence specificity and topology are not critical for Msdps2 molecules for its interaction.

Additionally, we performed EMSA of DNA-MsDps2 complexes with fluorophore-labelled 50 bp DNA on a Native PolyAcrylamide Gel Electrophoresis (PAGE) gel. This experiment suggested that with increased protein concentration (4 μM and 8 μM), fuzzy bands were observed in the middle of the gel, unlike a prominent free DNA band was observed in the control DNA lane (Figure 1E). However, when the protein concentration was increased dramatically (12 μM), the significantly large MsDps2-DNA complex could be formed, which appeared to be entirely stuck in the well of the Native PAGE gel. This observation suggested that at lower concentrations of MsDps2, a varied number of MsDps2 molecules bind to the 50 bp DNA, resulting in a blurry band migrating corresponding distances in the Native PAGE gel (Figure 1E).

Furthermore, we cross-validated the binding affinity of MsDps2 to DNA using MicroScale Thermophoresis (MST). We calculated the dissociation constant (K_D_) of 48 nM using the MST profile (Figure. 1F). These biochemical and biophysical studies suggested that MsDps2 binds with DNA. To dissect the architecture of the DNA-MsDps2 complex, we turned to high-resolution TEM imaging of the DNA-MsDps2 complex.

### Negative stained TEM analysis of Msdps2 binding on different DNA

MsDps2 does not carry a canonical DNA binding site, making it difficult to predict the binding conformations on DNA. To decipher the DNA binding interactions of MsDps2, we carried out room-temperature negative stained Transmission Electron Microscopy (NS-TEM) studies with different types of DNA. Initially, we visualized the MsDps2 alone by NS-TEM (Supplementary Figure 2A), which showed spherical, monodispersed oligomers. The reference-free 2D class averages of MsDps2 indicated a smooth surface of MsDps2 oligomer (Supplementary Figure 2B). However, when we incubated MsDps2 with 200 bp, 3 kb plasmid, and Holliday junction DNA, we observed complexes with different numbers of MsDps2 (Figure 2A-D and Figure Supplemental 2C-D). We again performed reference-free 2D classification of MsDps2 with 200 bp DNA and Holliday junction DNA for better understanding. We observed two, three, or four MsDps2 molecules bind to DNA, where lung-shaped DNA-MsDps2 complexes appeared predominantly (Figure 2E-G). These 2D class averages clearly showed an extra density between two MsDps2 molecules, which tightly holds the two MsDps2 molecules. However, most of the class averages were highly heterogeneous and adopt various conformations due to the conformational flexibility of DNA (Figure 2F).

**Figure 2:**
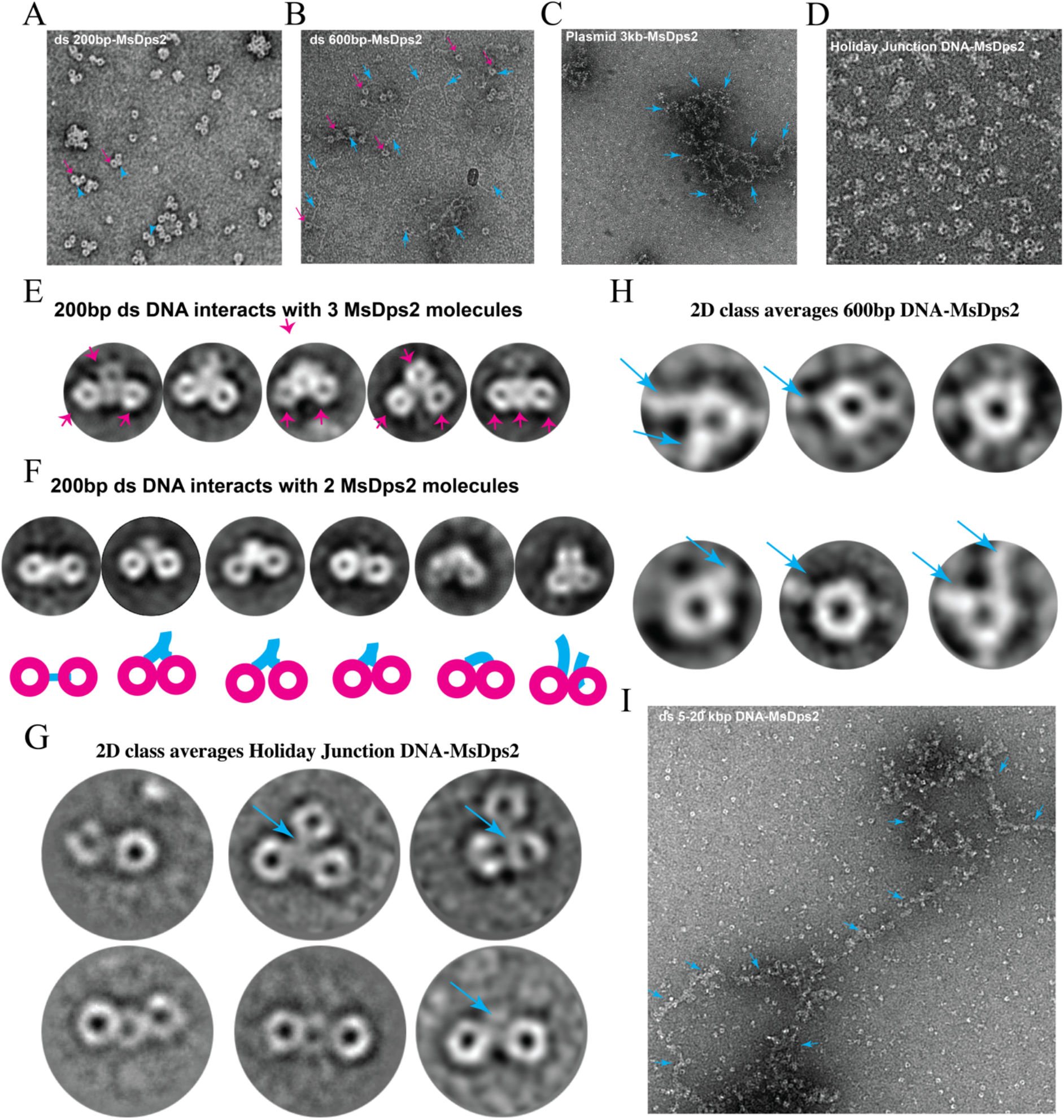
Negative staining of MsDps2-DNA: (A) Raw Micrograph of MsDps2 incubated with 200bp dsDNA. (B) Raw Micrograph of MsDps2 with 600bp dsDNA. (C) Raw Micrograph of MsDps2 with 3kb plasmid DNA. (D) Raw Micrograph of MsDps2 with Holiday Junction DNA. (E-F) Reference-free 2D class averages of MsDps2 with 200bp. (E) shows three MsDps2 molecules are attached to DNA whereas (F) shows two of the MsDps2 particles are attached DNA.The schematic diagram showed different conformation of MsDps2-DNA complex where pink colour circle denotes MsDps2 and blue colour denotes DNA. (G) Reference-free 2D class averages of MsDps2 with Holiday junction DNA. (H) Multi-site attachment of DNA in MsDps2. (I) Representative 2D classification of MsDps2 with Holiday junction DNA. The DNA was denoted with blue arrow.

Upon incubating with 600 bp DNA, we observed a thread-like extended architecture in the NS-TEM micrographs (Figure 2B). This extended thread-like architecture was completely missing in the DNA-free MsDps2 micrographs, indicating that DNA facilitates the long threadlike assemblies. Most interestingly, DNA-MsDps2 formed a bead-on-strings-like architecture where DNA acts as a string and MsDps2 as beads. The bead-on-strings-like DNA-protein complexes were selected to perform 2D class averages. We observed that each MsDps2 molecule surprisingly showed more than one binding site for DNA (Figure 2H). However, for longer DNA substrates (5 kbp-21 kbp), we did not observe these types of bead-on-strings-like assemblies. Negative stain TEM data suggest that MsDps2 interacted with 21kbp DNA and formed small clusters and finally small clusters interacted with other clusters to form large assembly. As a result, we could not solve the binding arrangement from NS-TEM images (Figure 2I and Figure Supplemental 2 E). Our 2D class averages of NS-TEM images showed that MsDps2 could interact with all types of DNA (linear DNA, plasmid DNA and Holliday junction DNA). In all 2D class average structures of TEM images – only MsDps2 or Mspds2-DNA complexes – MsDps2 appeared as uniform spherical particles, indicating that DNA binding does not alter the overall morphology of the dodecamer core. Instead, DNA was attached near the periphery of MsDps2, as shown in NS-TEM images.

NS-TEM micrographs and 2D class averages revealed crucial binding modes of MsDps2 with DNA which depict the underlined molecular interactions of the MsDps2-DNA complexes. However, to obtain the high-resolution structure of the complex at near physiological condition, we implemented the single-particle cryo-Electron Microscopy (Cryo-EM).

### Cryo-Electron Microscopy (cryo-EM)-based structural studies of MsDps2-DNA

MsDps2 is a member of the ferritin superfamily comprising multi-subunit cage-like proteins with a hollow interior (Roy *et al*., 2008). Structural elucidation of the MsDps2 binding on DNA is essential to understand how the DNA is condensed and protected by MsDps2. Our current biochemical and negative staining TEM data suggest that MsDps2 binds DNA via multiple binding sites. We carried out single-particle cryo-Electron Microscopy (cryo-EM) to probe interacting residues, which are crucial for the binding with MsDps2. The cryomicrographs indicated that MsDps2-DNA particles appear as beads-on-the-thread-like structures. Small rod-like densities were detectable on MsDps2 molecules in most 2D class averages (Figure 3A, Supplementary figure 3). Notably, this rod-like density along the MsDps2 was completely absent in the cryo-EM 2D class averages of MsDps2 alone. Altogether, these results strongly suggested that the rod-like densities indicate DNA associated with MsDps2 molecules. These results motivated us to determine the cryo-EM structure of the MsDps2-DNA macromolecular complex. Initially, we performed two-three rounds of reference-free 2D classification on the cryo-EM data to remove disordered, beam-damaged, and ice-contaminant particles. The resulting clean dataset was then used to perform heterogeneous refinement with two different particle diameters (136 Å and 200 Å) to isolate and separate out the structural heterogeneity of any MsDps2-DNA complexes.

**Figure 3:**
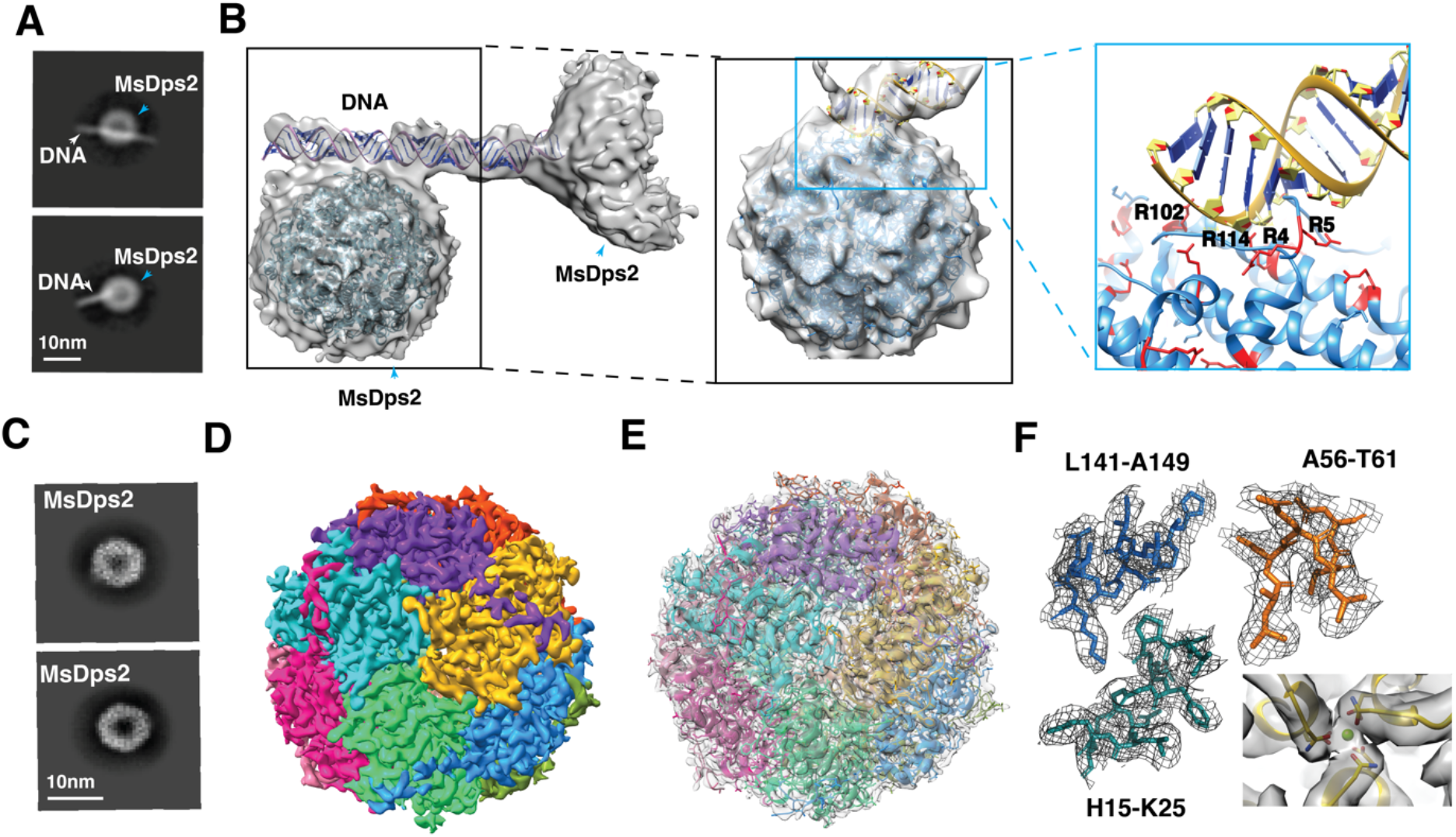
Cryo-EM map of MsDps2 DNA (600bp): (A) Representative reference free 2D class averages of MsDps2-DNA where DNA is denoted by white arrow head and MsDps2 is denoted by blue arrow head. (B) Cryo-EM map of MsDps2-DNA where DNA is attaching to two MsDps2 molecules and its docking with atomic model of MsDps2 and modelled DNA. Focused cryo-EM map of MsDps2-DNA as shown in black box. Enlarged view of MsDps2-DNA interaction region as shown in blue box. (C) Representative reference free 2D class averages of only MsDps2. (D) Cryo-EM map of MsDps2 with each protomer coloured with different colour. (E) Fitting of grey colour cryo-EM map of MsDps2 with atomic model generated by phenix refinement. (F) Fitting of side chains at different regions of wild type MsDps2 shows proper fitting of amino acid residues in the cryo-EM map.The Magnesium ion was docked in the cryo-EM density was shown in green colour ball.

Interestingly, all the 3D classes with particle diameter 136 Å showed extra DNA density tangential to MsDps2, which indicates that most of the MsDps2 molecules were attached to DNA (Supplementary Figure 3). Using this 3D classification technique, we could separate different conformations of MsDps2 molecules bound with DNA. However, some classes of this 3D classification, like class 05 and class 10, showed a shorter DNA density near the periphery of MsDps2. However, when 3D classification was performed with larger particle diameter (200 Å), we could segregate different complexes such as a DNA molecule with either one or two MsDps2 molecules. Although, the significant population of particles falls into class 9, where the density of DNA was not visible when two MsDps2 molecules were present. MsDps2-DNA complex adopts various structures due to inherent flexibility of DNA. Furthermore, DNA does not wrap around the MsDps2 molecules like histone protein, which make this extremely challenging to resolve atomic resolution structure of MsDps2-DNA.

Finally, we selected a few 3D classes for the high-resolution structural characterization of MsDps2-DNA complexes. We determined the 3D structure of the MsDps2-DNA complex at a resolution of 6.4 Å (Figure 3B). We speculated that due to the inherent structural flexibility of DNA molecules, MsDps2-DNA complexes adopt many different conformations. This compromised our efforts to resolve the atomic resolution structure of the MsDps2-DNA complex.

Furthermore, we performed the focused refinement of only MsDps2 that are part of DNA-MsDps2 complexes by imposing T-symmetry. This successfully resolved the 3.2 Å resolution cryo-EM structure of MsDps2 (Figure 3D). This indicates MsDps2 adopts a stable structure, even it interacts with DNA. The MsDps2 map at 3.2 Å was suitable for model building, and an atomic model was generated to identify critical residues for DNA-protein interactions (Figure 3E). Individual regions of the cryo-EM map and fitted model were analyzed separately to determine the overall quality of map. Many side chains, like H15-K23, A56-T61 and L141-A149, were well-fitted into the cryo-EM map (Figure 3F). Most interestingly, Mg^+2^ ions were also visible on our map. The combination of high-resolution MsDps2 and low-resolution entire complex MsDps2-DNA guided us to manually dock the atomic model into the complex. Thus, the atomic model built from our cryo-EM structure of MsDps2 core (PDB - 2Z90) and the computationally generated DNA structure were manually fit into the cryo-EM density map. The overall fitting of the atomic model strongly supports the moderate-resolution cryo-EM map. Further, the modelled structure of the DNA and atomic model of MsDps2 structure could provide sufficient information to perform site-directed mutation studies. Based on our docking results of the cryo-EM map of MsDps2-DNA, we predicted that amino acid residues Arg4, Arg5, Arg102, and Arg114 are responsible for DNA binding.

For comparing the structural changes of MsDps2 residues after incubating with DNA, we have also performed the 3D reconstruction for only MsDps2 (Supplementary figure 4). We achieved the atomic resolution structure of MsDps2 core of 2.2-3.0 Å resolution (local resolution calculation) and 2.94 Å (global resolution calculation) with T symmetry (Supplementary figure 4D) at 0.143 FSC (Supplementary figure 4E). There was no change in the conformation of MsDps2 as shown in biophysical experiment. However, we could see only minor changes in the orientation of side chains of those residues which might be involved in DNA binding interaction.

### Determining the interacting amino acids residues between DNA and MsDps2

As mentioned earlier, one of our aims was to identify the exact amino acid residues responsible for DNA binding. We were able to see the DNA density near the N terminal in our cryo-EM map of MsDps2-DNA complex. Moreover, the previous study suggested that N-terminal residues of MsDps2 might play a vital role in DNA binding (Roy *et al*., 2008). Therefore, we decided to truncate the N-terminal flexible loop to identify the DNA binding affinity of resultant MsDps2, which we call ΔN_15_MsDps2 (i.e., first 15 amino acids at the N-terminal deleted). Similar to wild-type protein, we puurified DNA-free ΔN_15_MsDps2 for DNA binding studies (Supplemental figure 6 A-B). We also performed EMSA to identify the DNA binding with ΔN_15_MsDps2. We noticed that DNA could interact with ΔN_15_MsDps2 (Figure 4A-B). We performed negative staining TEM studies to confirm our findings to observe ΔN_15_MsDps2-DNA interactions with 21 kb of DNA (Figure 4C). Our negative staining data suggested that ΔN_15_MsDps2 can interact with DNA. It also showed that the N-terminal deletion did not affect the oligomerization and formation of ΔN_15_MsDps2 dodecamer. This biochemical and negative staining data strongly suggested that other amino acid residues are also responsible for DNA binding apart from the N-terminal region.

**Figure 4:**
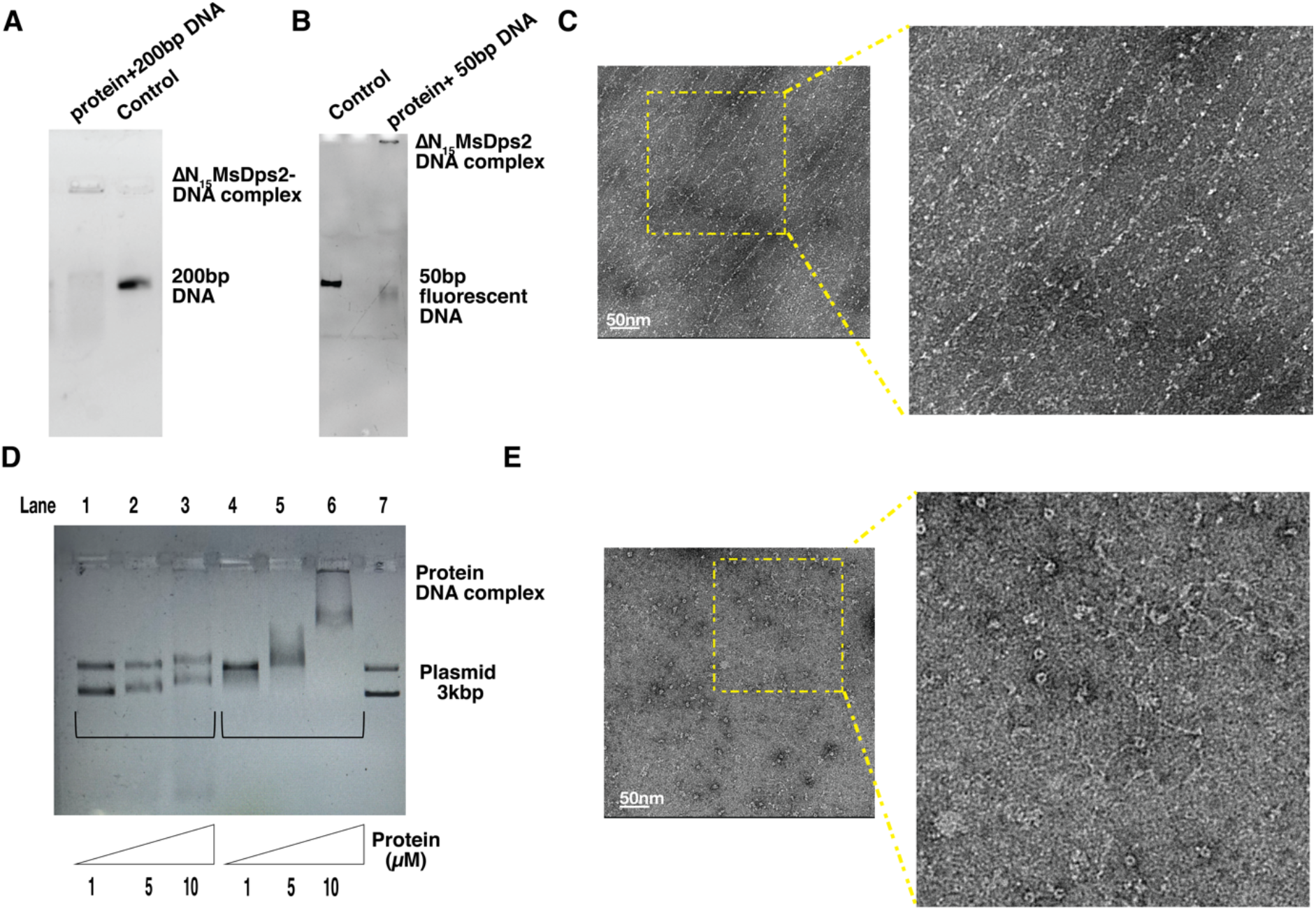
Biophysical and biochemical analysis of ΔN_15_MsDps2 with DNA and mutant MsDps2: (A) Electrophoretic mobility shift assay showing binding of ΔN_15_MsDps2 with 200bp DNA. (B) Native PAGE showing binding of ΔN_15_MsDps2 with 50bp fluorescent DNA. (C) Negative staining of ΔN_15_MsDps2 with 21kbp DNA. (D) Electrophoretic mobility shift assay. Lane 7 is only 3kb plasmid DNA. Lane 1-3 corresponds to 3kb plasmid DNA incubated with mutant MsDps2 with increasing protein concentration per well whereas Lane 4-6 corresponds to 3kb plasmid DNA incubated with wild type MsDps2 with increasing protein concentration per well. (E) Negative staining of ΔN_15_MsDps2 with 21kbp DNA.

Based on our docking results of cryo-EM map of MsDps2-DNA, the predicted amino acid residues Arg4, Arg5, Arg102, and Arg114 were mutated to glutamic acid (Arg4Glu, Arg5Glu, Arg102Glu, Arg114Glu) and purified the resultant MsDps2 variant. This variant of MsDps2 was examined by EMSA and negative staining imaging to study its DNA binding capability (Figure 4D-E). In both experiments, we noticed that DNA binding affinity was severely impaired. EMSA studies showed that DNA shift was negligibly low. Similarly, TEM data showed very few MsDps2 molecules bound on DNA, while a significant fraction of unbound mutated MsDps2 molecules were visible in the background (Figure 4E). Together, these biochemical and biophysical studies suggested that Arg4, Arg5, Arg102, and Arg114 play a vital role in DNA binding but not completely abolishing the DNA binding property of MsDps2. To further identify the second binding site of MsDps2 molecules, another high-resolution structural map of either ΔN_15_MsDps2 or mutated MsDps2 is required. We selected ΔN_15_MsDps2 for delineating the remaining amino acid residues that interact with DNA.

### Cryo-EM-based structural studies of ΔN_15_MsDps2-DNA

We performed single particle cryo-EM of ΔN_15_MsDps2-DNA complex to identify the DNA binding residues of MsDps2 in the absence of the N-terminal domain (Supplementary Figure 7). We used 50 bp DNA instead of 600 bp DNA to reduce the inherent flexibility of DNA for the cryo-EM structural characterization of ΔN_15_MsDps2 interaction. Surprisingly, we could see an extension near ΔN_15_MsDps2, which is clearly visible in reference-free 2D class averages (Supplementary Figure 7). We also calculated a 3D structure of ΔN_15_MsDps2-DNA, which forms a “cherry hanging from a branch” like structure (Supplementary Figure 9A). The branch was the DNA, whereas ΔN_15_MsDps2 adopts a cherry-like structure. The 3D structure of the cryo-EM map was determined at a resolution of ~6.6 Å owing to the flexible nature of the large complex (Figure 5A). The local resolution plot of ΔN_15_MsDps2-DNA shows that the resolution is ranged between 4.8-12 Å (Supplementary Figure 9B). The focused refinement of ΔN_15_MsDps2-DNA core complex results in a high-resolution structure of ΔN_15_MsDps2 at 3.4 Å by imposing T-symmetry (Figure 5D). We generated an atomic model by performing Phenix refinement and fitted it into cryo-EM density (Figure 5E). The side chain fitting of some residues is shown in Figure 5F. The local resolution plot of ΔN_15_MsDps2 showed that the core was resolved between a resolution of 2.8-3.8 Å (Supplementary Figure 9C-D). The atomic model of ΔN_15_MsDps2 and DNA were fitted into the cryo-EM density map, which assisted us in identifying key amino acid residues responsible for the DNA-protein interaction (Figure 5B). We found that the Arg114, Arg102, Arg84, Arg124, and Arg 123 residues of ΔN_15_MsDps2 interact with DNA (Figure 5C). We performed Site Directed Mutagenesis of the amino acids to validate the DNA binding amino acids. However, resultant mutants could not express, and we could not perform any DNA binding assay.

**Figure 5:**
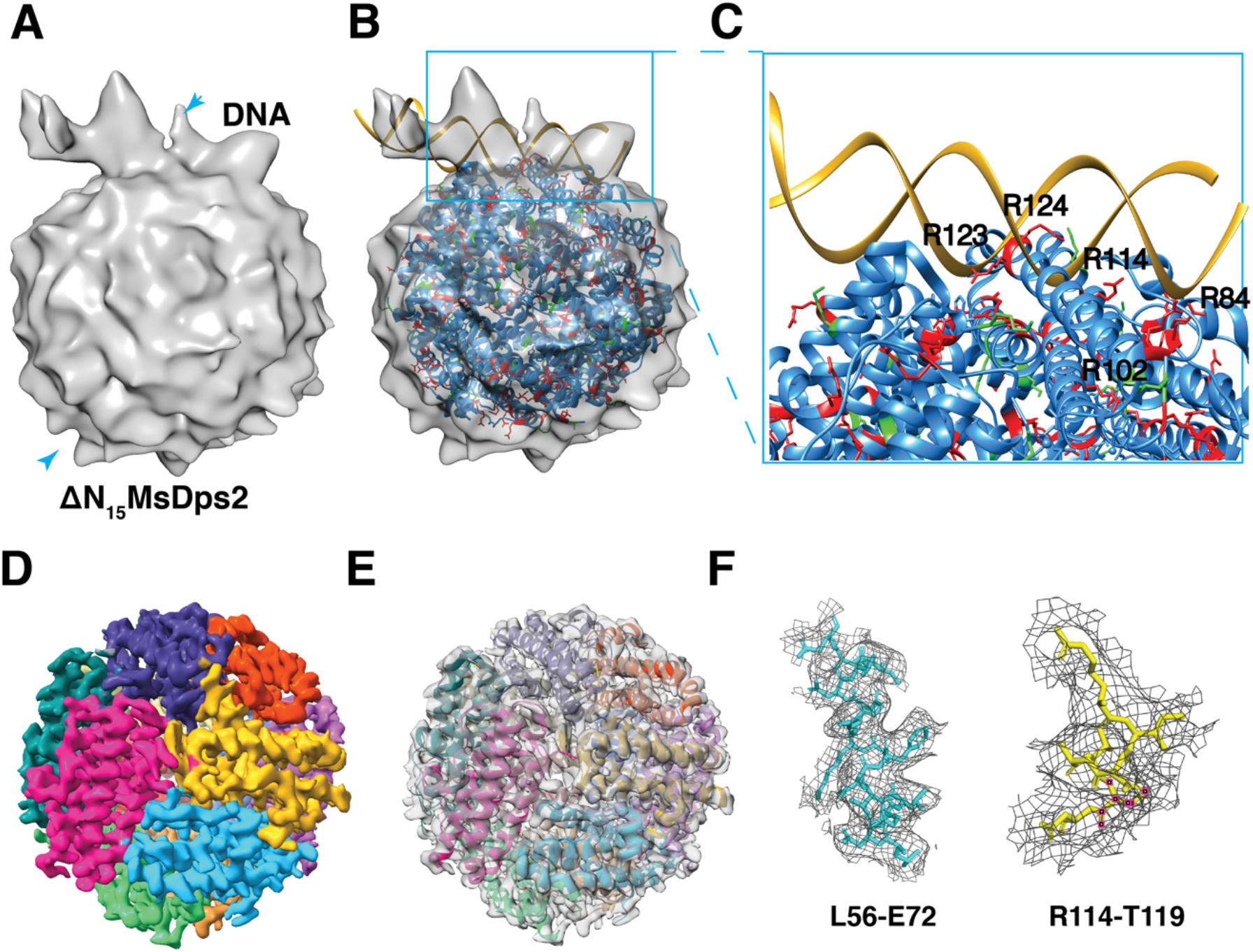
Cryo-EM map of ΔN_15_MsDps2 DNA (50bp): (A) Cryo-EM map of ΔN_15_MsDps2-DNA where DNA is attached to ΔN_15_MsDps2 (B) Docking with atomic model of ΔN_15_MsDps2 and modelled DNA of cryo-EM map. (C) Enlarge view of interacting residues of ΔN_15_MsDps2 with DNA (D) Cryo-EM map of ΔN_15_MsDps2 with each protomer coloured with different colour. (E) Fitting of grey colour cryo-EM map of ΔN_15_MsDps2 with atomic model generated by phenix refinement. (F) Fitting of side chains at different regions of ΔN_15_MsDps2 shows proper fitting of amino acid residues in the cryo-EM map.

### Single Molecule Imaging

Further, we wanted to test the DNA compaction abilities of WT MsDps2, ΔN_15_MsDps2, and mutant MsDps2 using real-time visualization at the single-molecule level. We performed singl-moleule fluorescence imaging of MsDps2 interactions with DNA. These experiments are similar to the ones published by Vtyurina et al. 2016 (Vtyurina *et al*., 2016). For this, we immobilized a 25-kilobasepair double-stranded DNA modified with single biotin at one of its ends on a PEG-passivated glass surface. We visualized DNA molecules under Total Internal Reflection Fluorescence (TIRF) microscope using an intercalating dye called Sytox Orange. We observed immobilized DNA molecules fluctuating around the immobilized position providing a diffusive fluorescence signal as seen in WT MsDps2 at zero seconds time point (Figure 6). Next, we introduced 1 μM MsDps2 into the flow cell, where DNA molecules were immobilized. We observed DNA molecules tightly compacting within 20 s appearing as fluorescent spots without any fluctuations. We then tested the ability of ΔN_15_MsDps2 to compact DNA using the same assay. When we flowed in a 1 μM of ΔN_15_MsDps2 onto immobilized DNA, we did not observe any considerable change in the fluctuations of tethered DNA molecules. However, when the ΔN_15_MsDps2 concentration was 2 μM, we observed DNA compaction similar to the wild-type protein.

**Figure 6:**
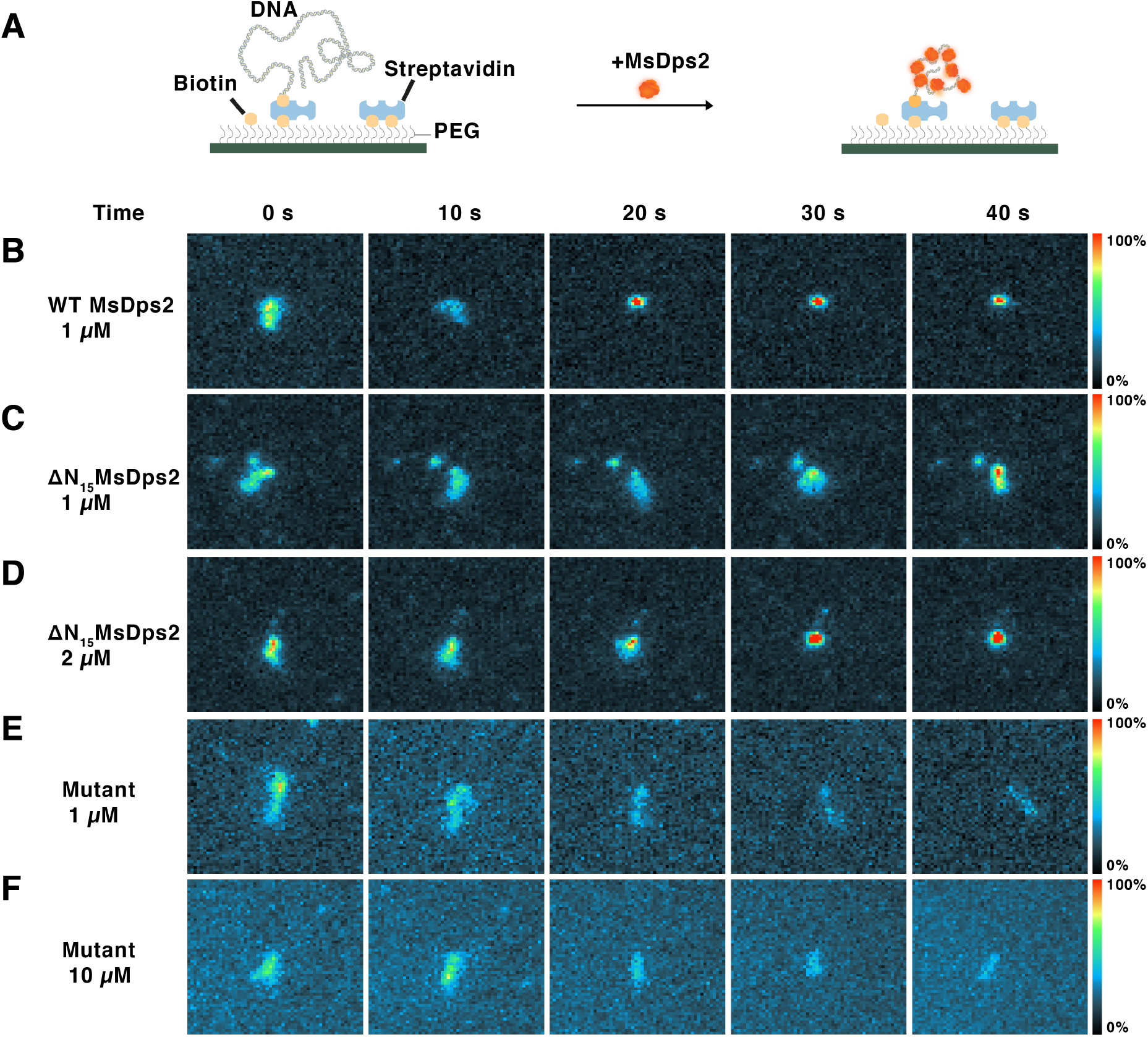
Single-molecule visualization of DNA compaction by MsDps2: A) Left – Schematic diagram showing immobilized DNA on a PEG-passivated surface via biotin-streptavidin interactions. Right – Schematic showing compacted DNA by MsDps2 (red). B) Series of images showing single fluorescent DNA molecule under TIRF microscope undergoing compaction by wild-type MsDps2. In first two images, DNA is non-compacted as seen by diffusive fluorescence signal and appeared as fully compacted from third image (i.e., 20 s) onwards. C) DNA did not compact at 1 μM ΔN_15_MsDps2. D) At 1 μM ΔN_15_MsDps2, DNA is seen compacted from 30 s onwards. E) and F) Mutant Dps2 did not show any compaction even at a concentration of 10 μM.

Similarly, we wanted to perform single molecule imaging experiments of the mutated MsDps2 (Arg4Glu, Arg5Glu, Arg102Glu, Arg114Glu) to understand the affinity of DNA with mutated MsDps2. We examined a ten times higher concentration of mutated MsDps2 compared to the wild type for single molecule imaging experiments. The results showed no detectable difference between the free DNA and DNA incubated mutated MsDps2 in terms of DNA fluctuations, which indicating mutated MsDps2 does not compact DNA. These singlemolecule imaging results strongly corroborate our electrophoretic mobility shift assay, TEM, and cryo-EM data (Figure 6).

### Proposed model of MsDps2 compacting DNA

Our current data from our single particle cryo-EM and single molecule imaging confirm that MsDps2 interacts with DNA and high concentration of MsDps2 protein is required for DNA compaction. Thus, we visualized the situation where large genomic DNA was incubated with very high concentration of protein at cryogenic conditions. Surprisingly, we noticed that MsDps2 molecules were formed a 2D crystal lattice after incubating with genomic DNA (Figure 7A). We observed that one MsDps2 molecule was surrounded by six MsDps2 molecules in the hexagonal lattice structure as shown in the reference free 2D classes (Figure 7B). However, we were unable to visualize the DNA density in between MsDps2 molecules. This might be possible that flexible DNA was averaged out during image reconstruction. Cryo-ET (cryo-electron tomography) will be an alternative procedure to resolve this type of 2D lattice arrangement as shown in previous study (Kamyshinsky *et al*., 2019). Further, we generated the 3D reconstruction of hexagonal lattice (Figure 7C). Interestingly, we observed that a central density connected with three MsDps2 molecules in 2D class averages after reducing the particle diameter. This central density within three MsDps2 molecules might be the DNA which is perpendicularly associated with MsDps2 molecules (Figure 7D). Based on this observation, we proposed a model of how MsDps2 compacts DNA by producing a 2D crystal lattice, with DNA lying in each interspatial space between the molecules of MsDps2. We have also highlighted that crucial arginine residues required for DNA-MsDps2 interaction identified from the cryo-EM structure of the MsDps2-DNA complex as shown in Figure 7E.

**Figure 7:**
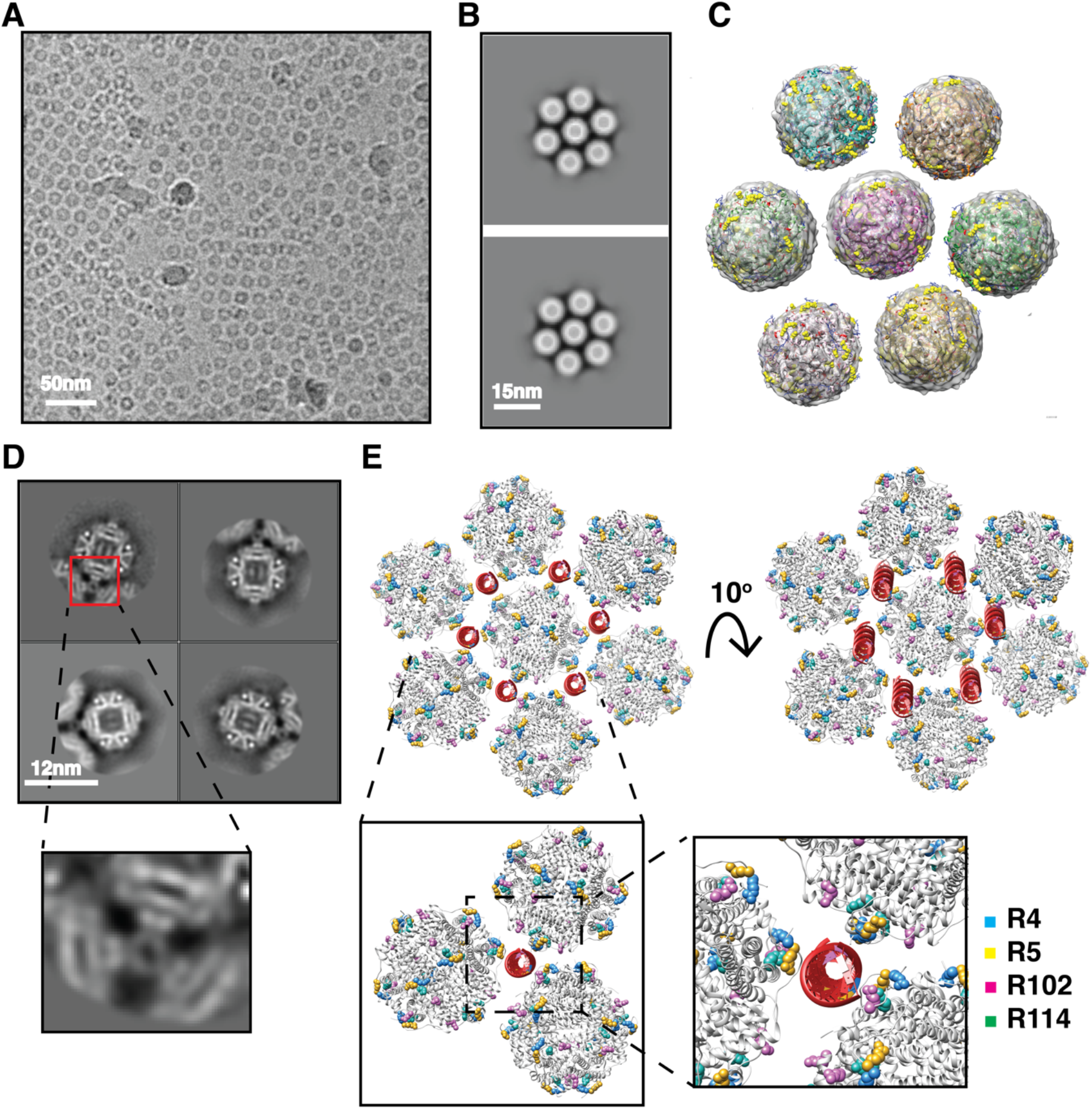
Proposed model for the mechanism of DNA compaction with MsDps2: A) The Representative cryo-EM Raw micrograph showing lattice formation. B) The representative reference free 2D class averages of lattice where one MsDps2 molecule is surrounded by six MsDps2 molecules. C) The docking of MsDps2 atomic model into the cryo-EM 3D map of MsDps2 lattice. D)The representative reference free 2D class averages focusing the three molecules of MsDps2 with central density corresponds to DNA and highlighted in red box. E) The proposed model for mechanism of DNA compaction by MsDps2 where DNA is residing in the interspatial space between three MsDps2 molecules present in hexagonal lattice. The important residues R4,R5,R102 and R114 of each MsDps2 (grey in colour) highlighted in blue, yellow, pink and green colour respectively are interacting with DNA (red in colour).

## Discussion

MsDps2 is a DNA-binding protein from *Mycobacterium smegmatis* starved cells, which protects its genome during hostile environmental conditions like starvation, oxidative stress, or antibiotic stress (Roy *et al*., 2008, Saraswathi *et al*., 2009). The structural insights of DNA binding motifs of Dps protein has been under investigation by many scientific groups (Tereshkin *et al*., 2019, Dadinova *et al*., 2021, Dubrovin *et al*., 2021). Few studies utilized AFM and negative staining TEM to visualize the DNA-Dps complexes. Based on these 2D images, three-dimensional structures were speculated (Dubrovin *et al*., 2021). Some studies showed that Dps can form a 2D array in the presence of DNA which was imaged using cryoelectron tomography to characterize the 3D structure of the Dps-DNA complex. However, none of these studies could determine the atomic resolution structure of Dps molecules during the interaction with DNA (Tereshkin *et al*., 2019, Dadinova *et al*., 2021, Dubrovin *et al*., 2021). In this study, we applied single particle cryo-EM to determine the high-resolution structures of the MsDps2-DNA complex. We reveal the key amino acid residues of Dps that play essential role in DNA binding and compaction activity.

Previous studies showed that Dps protein has histone-like properties (Drlica & Rouviere-Yaniv, 1987, Schmid, 1990). Furthermore, like histone-protein, Dps interact with DNA in sequence-independent manner (Berger, 1999). In line with these previous studies, our negative staining and EMSA data also demonstrated that MsDps2 could bind DNA sequences independently. Interestingly, negative staining and cryo-EM data strongly suggested that DNA interacts with MsDps2 at the periphery region, consistent with the observation of recent cryo-electron tomographic reconstruction of *E. coli* DNA (Kamyshinsky *et al*., 2019). However, as mentioned earlier, we intended to employ single particle analysis to obtain the high-resolution 3D structure of the MsDps2-DNA complex. To enable single-particle analysis, we required individually isolated particles. We examined comparatively diluted MsDps2-DNA samples where single MsDps2 molecules could be observed. Our single particle 3D reconstruction and docking studies showed that DNA binds to the N-terminal residues along with residues proximal to N-terminal in a tertiary form of MsDps2 molecules (Figure 3 and 5). In addition, due to relatively low protein-DNA concentration in cryo-EM and asymmetric reconstruction, DNA densities were not visible near the N-terminal/C-terminal regions of all protomers of the MsDps2 oligomer.

We showed that a low-resolution 3D structure of MsDps2-DNA was enough to perform rigid body docking and automatic docking of MsDps2 molecule and DNA separately. Furthermore, we employed focused refinement to resolve the atomic resolution 2.6-3.4 Å structure of only MsDps2, which is one of the first cryo-EM structures of Dps molecules from *Mycobacterium*. However, the atomic model of MsDps2 in our study plays a vital role in identifying amino acid residues that are responsible for DNA binding. Our biochemical and biophysical studies strongly support our structural studies and demonstrated that arginine residues of N-terminal (R4, R5) along with R102, R114 are predominantly responsible for DNA binding. Previous studies showed that N-terminal and C-terminal loops play an important role in DNA binding (Roy *et al*., 2007). However, our studies are the first structure-based study at near-native physiological conditions identifying amino acid residues located at various positions other than N- or C-terminal tail that are responsible for DNA binding.

Furthermore, single-molecule imaging analysis strongly supported structural and biochemical studies. Our study showed that except Arg4 and Arg5, the other two amino acids Arg102 and Arg114 were also responsible for DNA binding, which were proximal to the N terminal in tertiary structure of the Dps protein (Figure 3). Our single molecule imaging experiments showing DNA compaction by MsDps2 confirmed the phenotypes of MsDps2 variant and truncation observed under TEM, Cryo-EM imaging, and biochemical analysis studies.

In summary, we determined the cryo-EM structure of MsDps2 with DNA, where DNA binds to MsDps2 molecules near the N-terminal regions. Our study also revealed several arginine residues near N-terminal region are non-specifically interacting with DNA. Additionally, single-molecule imaging-based studies indicated that DNA compaction by WT MsDps2 showed a drastically reduced DNA compaction upon mutating the DNA interacting residues of MsDps2. We note that several other factors, like protein concentration, ionic strength, and pH of the buffer would play a vital role in DNA compaction, which will be explored in future studies.

## Acknowledgements

We acknowledge the Department of Biotechnology, Department of Science and Technology (DST) and Science, and Ministry of Human Resource Development (MHRD), India for funding and cryo-EM facility at IISc-Bangalore. We acknowledge DBT-BUILDER Program (BT/INF/22/SP22844/2017) and DST-FIST (SR/FST/LSII-039/2015) for National Cryo-EM facility at IISc, Bangalore. We acknowledge the financial support from the Ministry of Human Resource Development (MHRD) (Grant Number-STARS-1/171), and DBT (Grant No. BT/PR25580/BRB/10/1619/2017) for financial support. We thank DBT-IISc partnership program for TEM facility at Biological Sciences Division. We want to acknowledge the LC-MS facility and SPR facility at Biological Sciences Division for the acquisition of the mass spectrometry analysis and binding affinity characterization studies. I would like to acknowledge Prof. Dipankar Chatterji for providing us plasmid of MsDps2.I would like to thanks Dr. Sudhanshu Gautam for helping me in biophysical experiments. I would like to acknowledge Prathibhan KV and Sainath Polepalli for helping me in negative staining. I would like to acknowledge Sunanda Williams for critical evaluation of manuscript. I would like to acknowledge Nayanika Sengupta for evaluating the manuscript.

## Author credit statement

Priyanka Garg – Conceptualization, Investigation, Formal Analysis, Visualization, Writing – original draft and editing.

Thejas Satheesh – Perform single molecule experiments, Formal Analysis, Writing, Original draft, reviewing, editing.

Mahipal Ganji - Design single molecule experiments, Supervision, Formal Analysis, Writing, Original draft, reviewing, editing

Somnath Dutta – Conceptualization, Supervision, Formal Analysis, Resources, Visualization, Writing – original draft, reviewing, editing and funding.

## Data and materials availability

Cryo-EM electron density maps of Wild type MsDps2, Two MsDps2-DNA, MsDps2-DNA,Focused MsDps2, ΔN_15_MsDps2-DNA,Focused ΔN_15_MsDps2 have been deposited in the Electron Microscopy Databank and PDB. Wild Type MsDps2: PDB 8HX0, EMD-35070; Two MsDps2-DNA:EMD-35068; MsDps2-DNA:EMD-35072; Focused MsDps2:PDB 8HX1,EMD-35071; ΔN_15_MsDps2-DNA: EMD-35073;Focused ΔN_15_MsDps2: PDB 8HWZ, EMD-35069.All data needed to evaluate the conclusions in the paper are present in the paper and/or the Supplementary Materials.

