## Supplemental file1 for "Cryo-EM reveals the mechanism of DNA compaction by *Mycobacterium smegmatis* Dps2"

### Supplementary Figures

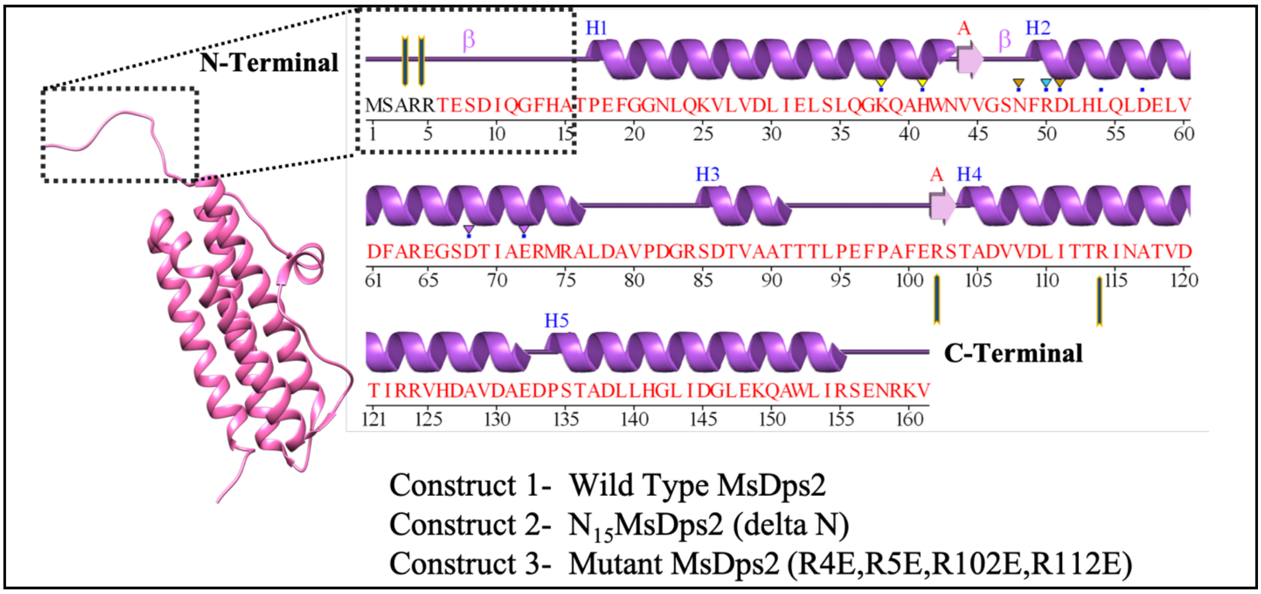


**Figure 1:Different constructs of MsDps2:** Residues deleted from construct 2 are highlighted with black box. Residues mutated in construct 3 are denoted by yellow arrow.

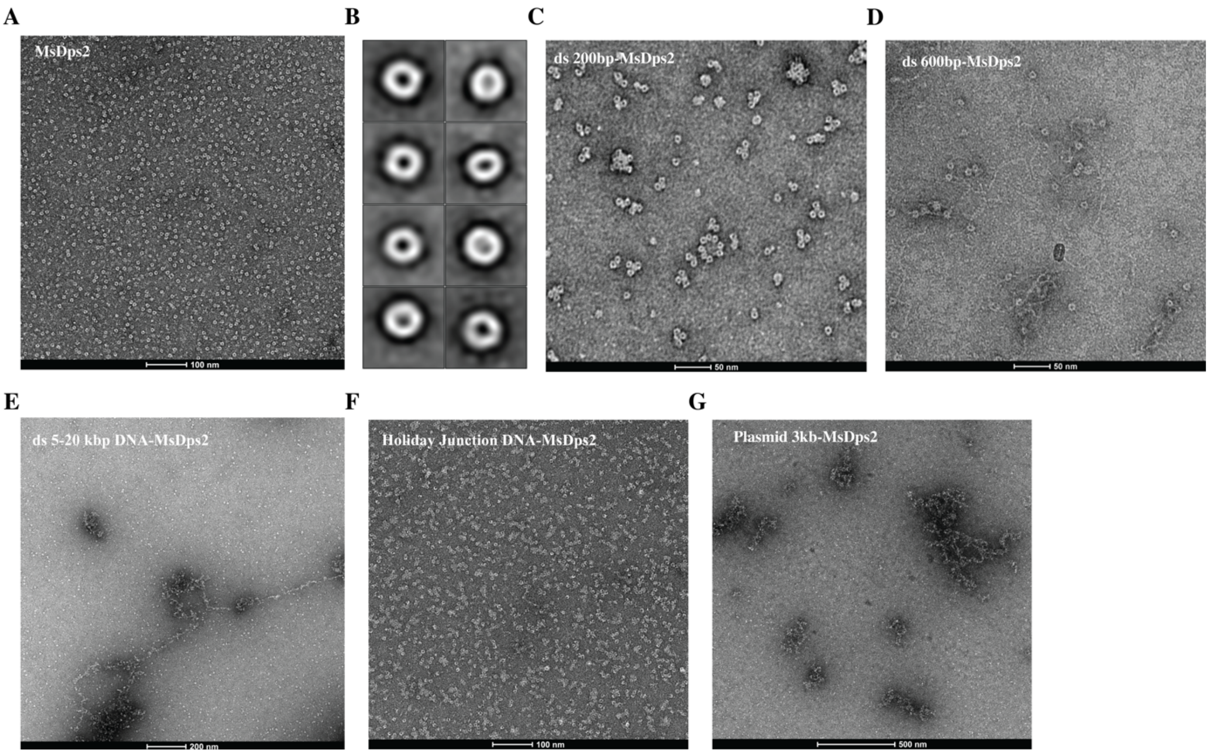

**Figure 2:Negative staining of MsDps2 and with different types of DNA:** (A) Raw Micrograph of only MsDps2. (B) Reference free 2D class averages of only MsDps2. (C) Raw Micrograph of MsDps2 incubated with 200bp dsDNA. (D) Raw Micrograph of MsDps2 with 600bp dsDNA. (E) Raw Micrograph of MsDps2 with 21kb dsDNA. (F) Raw Micrograph of MsDps2 with Holiday Junction DNA. (G) Raw Micrograph of MsDps2 with 3kb plasmid DNA.

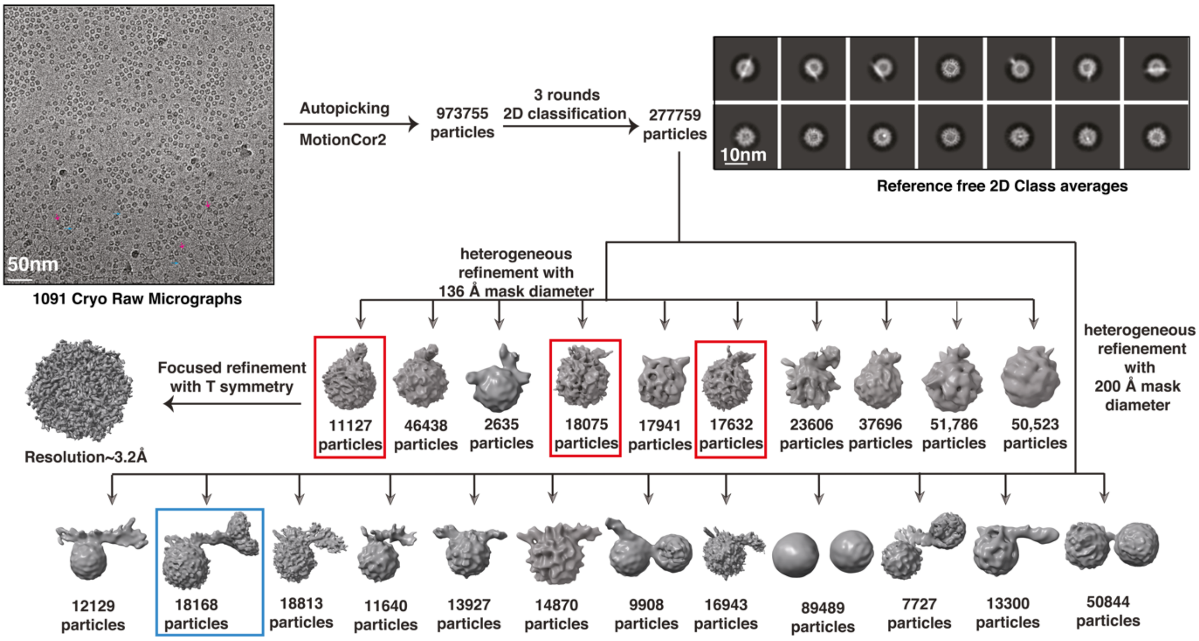


**Figure 3:Flow chart of Cryo-EM processing of MsDps2 with DNA 600bp.**

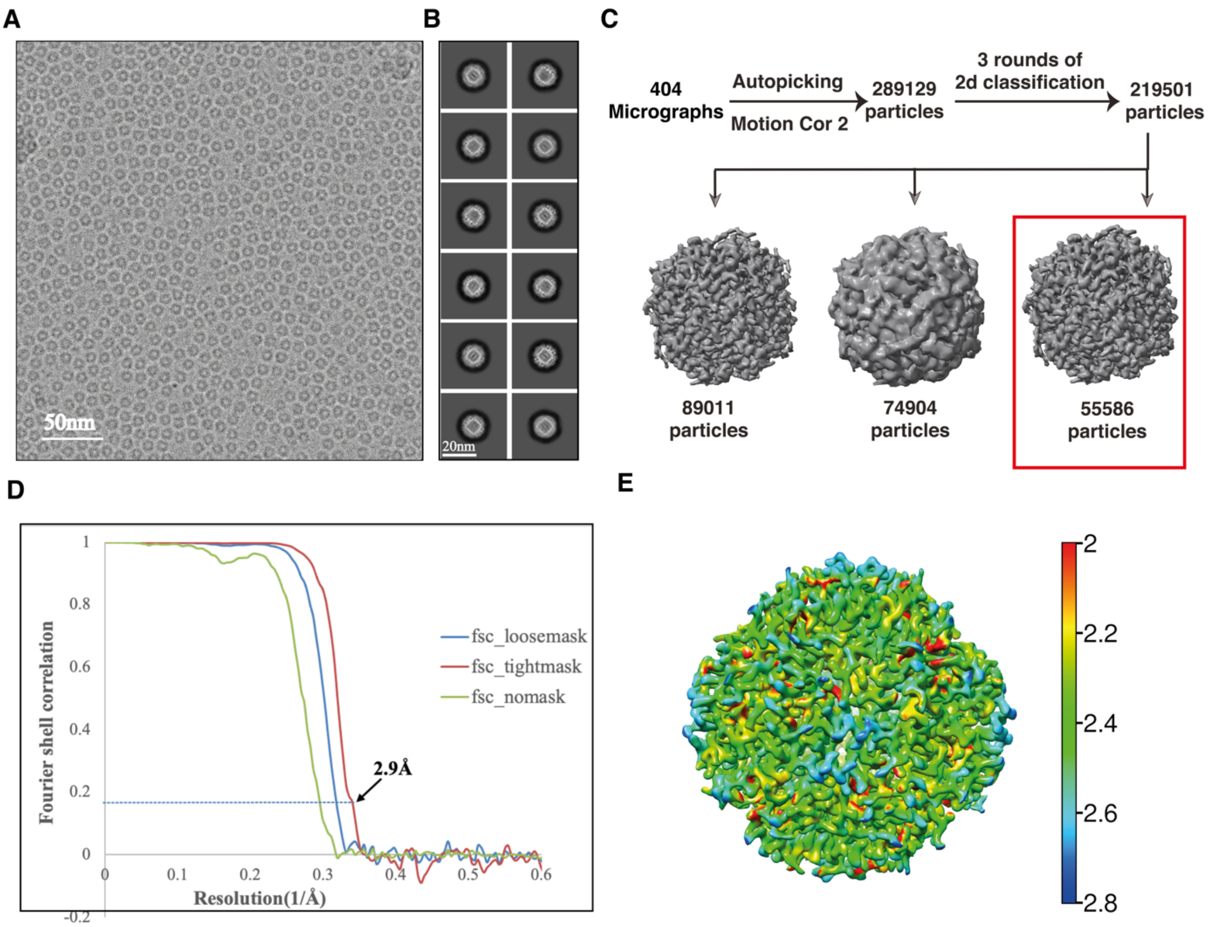


**Figure 4:Flow chart of Cryo-EM processing of MsDps2:** (A) Cryo-EM Raw Micrograph of only MsDps2. (B) Reference free 2D class averages of only MsDps2. (C) Flow chart of Cryo-EM processing of MsDps2. (D) FSC of only MsDps2 showing 2.9Å resolution. (E) Resmap of only MsDps2

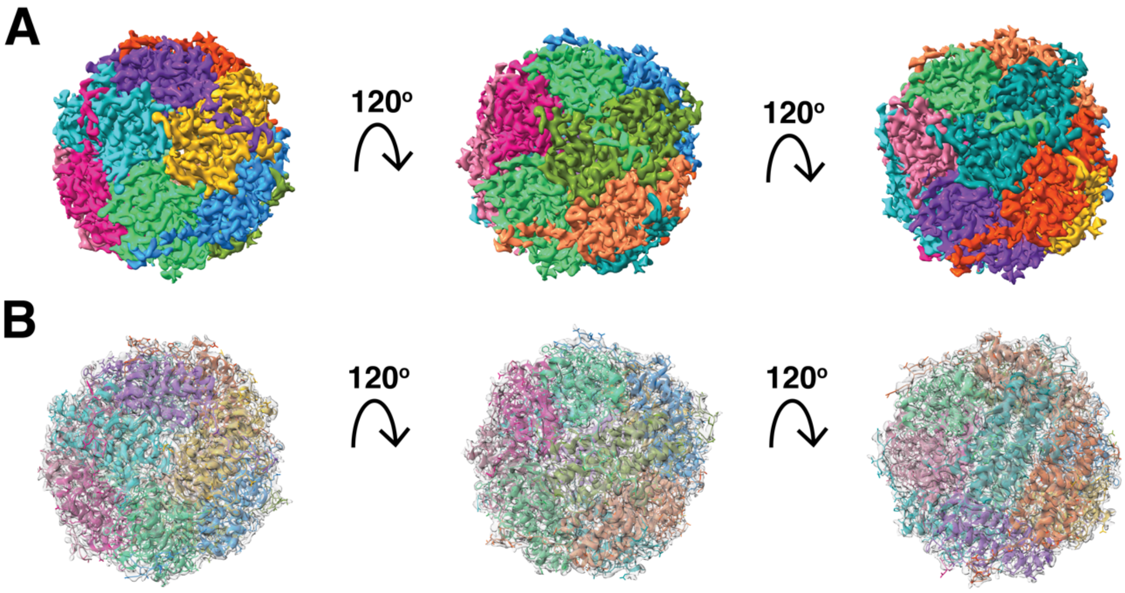


**Figure 5:Cryo-EM map of MsDps2:** (A) Different orientation of coloured map of MsDps2. (B) Different orientation of grey colour cryo-EM of MsDps2 fitted with atomic model generated by phenix refinement.

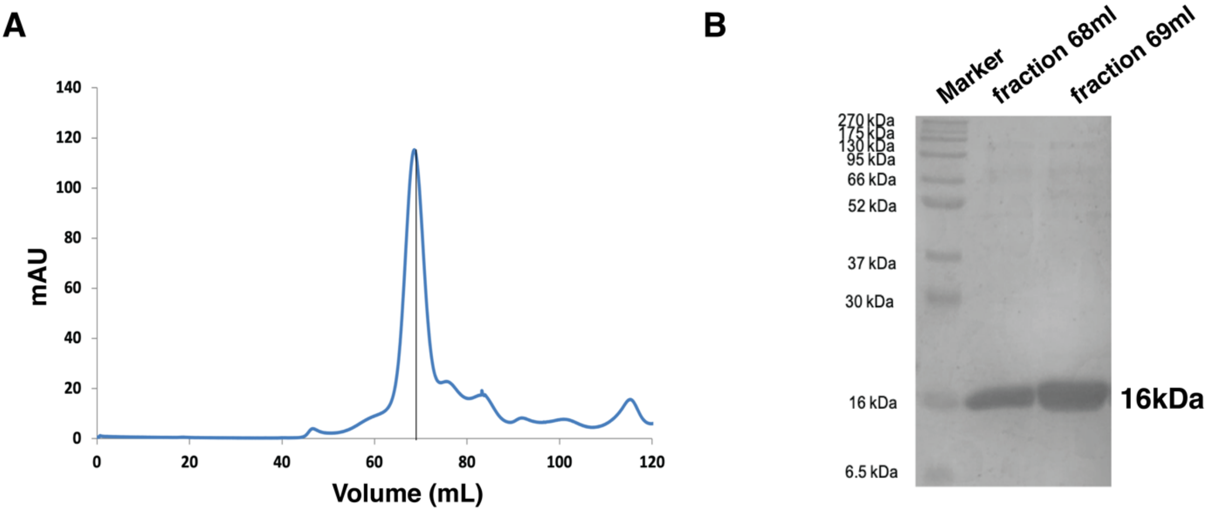


**Figure 6:Biophysical characterisation of $\Delta N_{15}$MsDps2 and its binding with DNA:** (A) shows the SEC profile for $\Delta N_{15}$MsDps2 that shows formation of a single peak at the expected molecular weight (198 kDa). (B) SDS PAGE analysis of $\Delta N_{15}$MsDps2 protein.

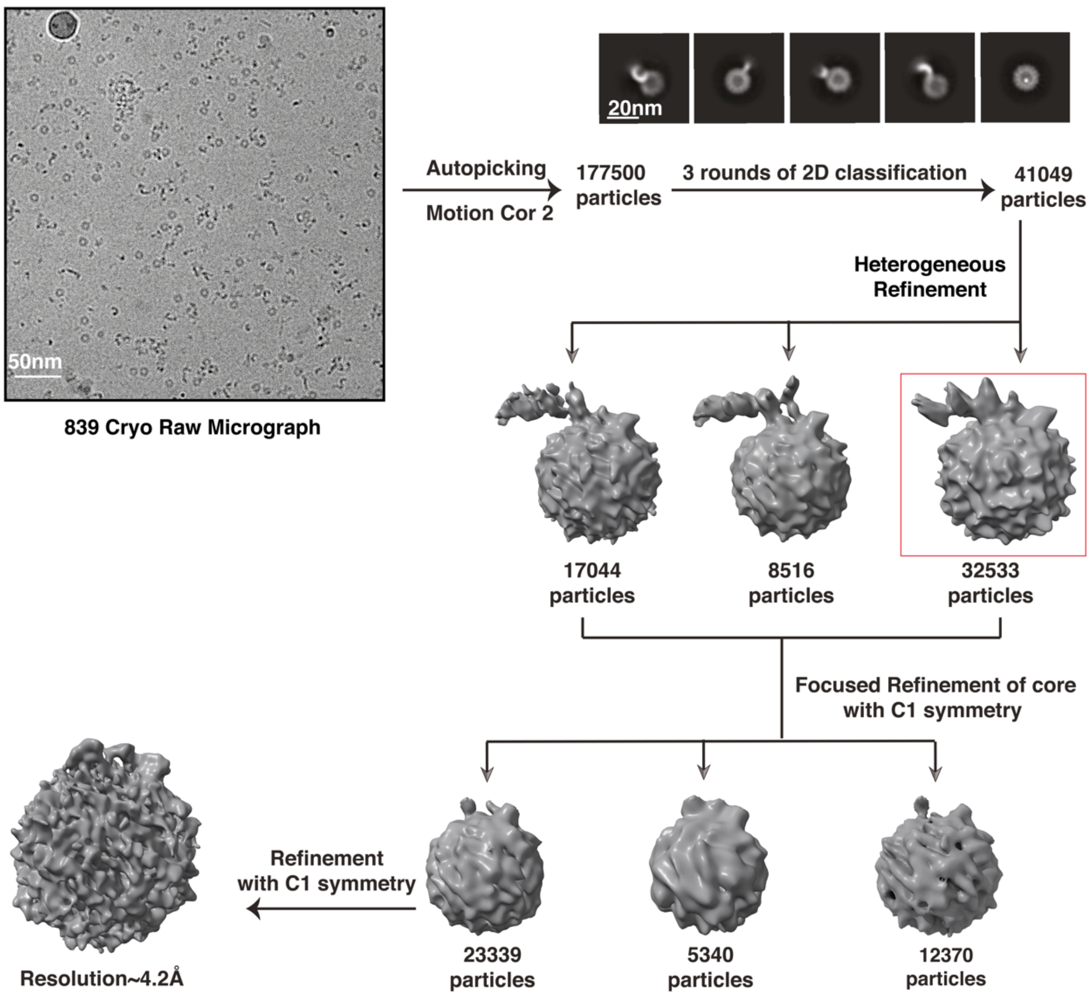


**Figure7: Flow chart of $\Delta N_{15}$MsDps2 with DNA.**

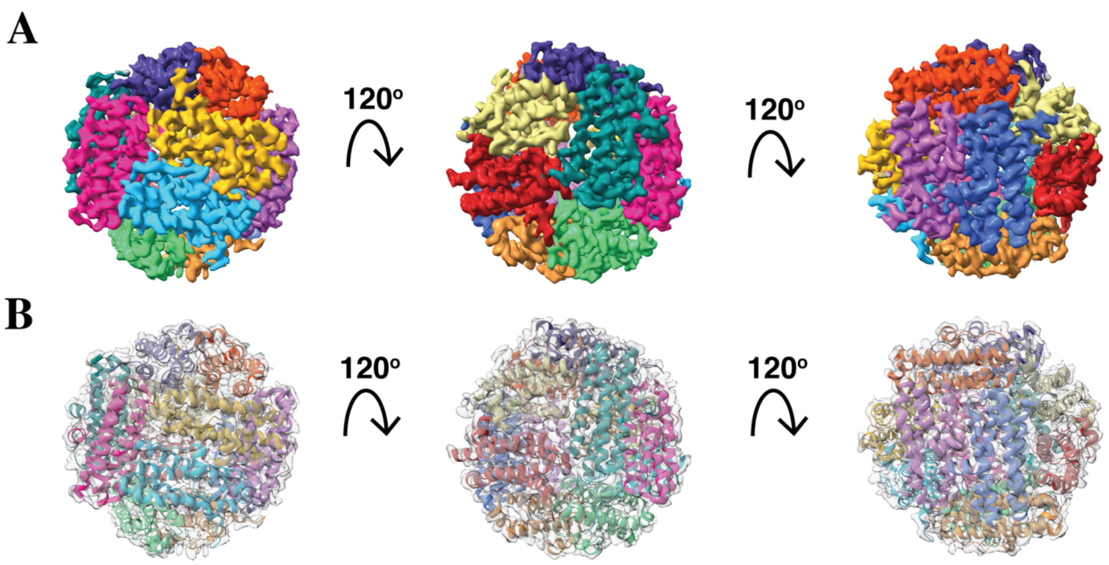

**Figure 8:Cryo-EM map of ΔN₁₅MsDps2:** (A) Different orientation of coloured map of only ΔN₁₅MsDps2. (B) different orientation of grey colour cryo-EM map of ΔN₁₅MsDps2 fitted with atomic model generated by phenix refinement.

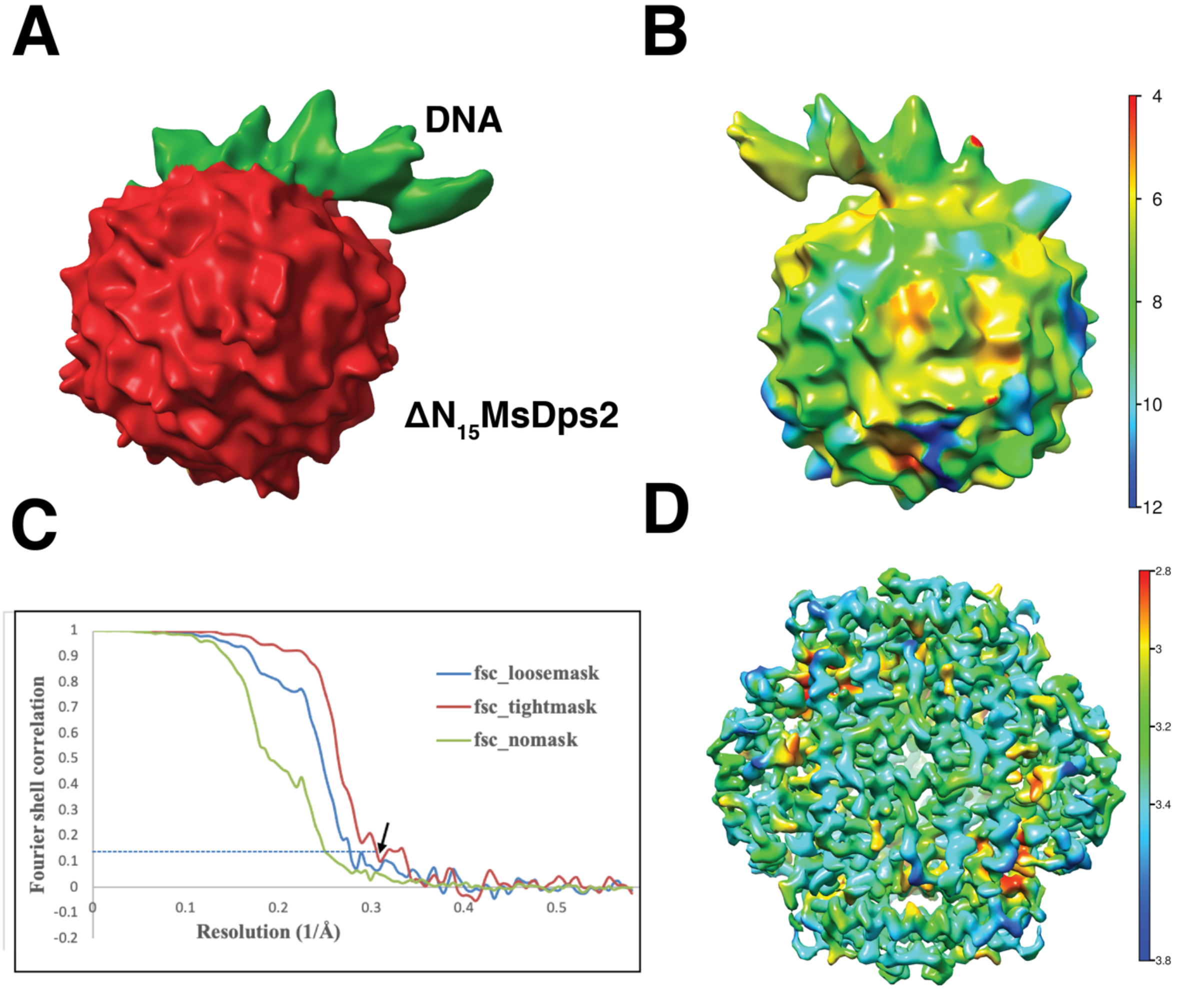


**Figure 9:** (A) Coloured cryo-EM map of ΔN₁₅MsDps2 with DNA where ΔN₁₅MsDps2 is red in colour and green colour corresponds to DNA. (B) Resmap of ΔN₁₅MsDps2 with DNA. (C) FSC of only cryo-EM of ΔN₁₅MsDps2. (D) Resmap of cryo-EM of ΔN₁₅MsDps2.

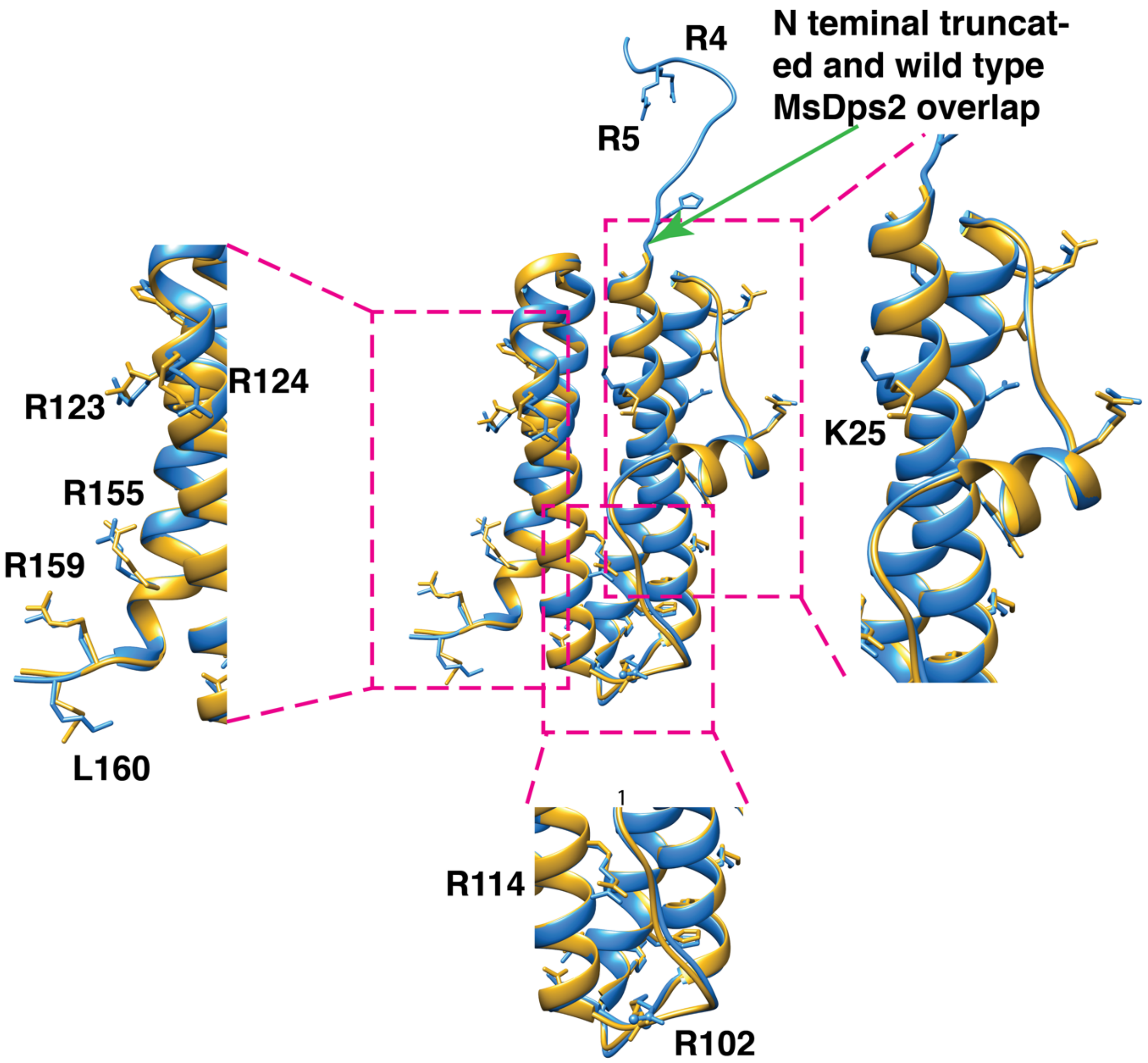


**Figure 10: Overlap of monomer model of $\Delta N_{15}$MsDps2 and Wild Type MsDps2**: Side chains of important residues required for DNA binding were highlighted.

**Table S1: List of plasmids and strains used in this study**

| **Strains and plasmids** | **Description** | **Antibiotics Selection** | **Reference** |
|---|---|---|---|
| *E. coli* DH5α | *E. coli* host strain for cloning purpose | Nil | Lab stock |

| | | | |
|---|---|---|---|
| *E. coli* BL21(DE3) | *E. coli* host strain for protein expression | Nil | Lab stock |
| *MsDps2_wt_pET21b* | *MsDps2* gene was cloned between NheI and HindIII, Expressed without tag. | Amp | Roy *et al.*,2008 |
| *ΔN₁₅MsDps2_pET21b* | 16-162aa PCR product of *MsDps2* gene was cloned between NheI and HindIII, to expressed *ΔN₁₅MsDps2* without tag. | Amp | This Study |
| *MsDps2_mutant_pET21b* | *MsDps2_wt_pET21b* plasmid was used to generate a point mutation at the 4$^{th}$, 5$^{th}$, 102$^{nd}$ and 114$^{th}$ position with the help of Site-directed mutagenesis (SDM) in four rounds of amplification. | Amp | This Study |

**Table S2 - List of primers used for different cloning**

| Primer Name | Sequence 5'→3' |
|---|---|
| DPS2-R2-3E-FP | *CTGGTCATGAGCGCAGAAGAGACTGAATCAGATATCCAA* |
| DPS2-R2-3E-RP | *TCCAACGCTAGCCATATGTATATCTCCTTCTTAAAGTTAAACAAAATTATTTC* |
| DPS2-R102-FP | *CCCCGCGTTCGAGGAGAGCACAGCCGATGTCGT* |
| DPS2- R102-RP | *AATTCCGGCAGCGTCGTGGTGG* |
| DPS2-R114-FP | *CGTGGCGTTGATCTCGGTGGTGATGAGGTC* |
| DPS2- R114-RP | *GACACCATCCGGCGCGTCC* |

| | |
|---|---|
| DPS2-DeltaN-FP | *ATATATGCTAGCGCCACACCGGAGTTCGGCGGC* |
| DPS2-DeltaN-RP | *ATATATAAGCTTTTAGACCTTCCTGTTCTCCGAGC* <br> *GGATCAGCCACG* |

**Table S3: Cryo-EM data acquisition, processing parameters and refinement statistics**

| **Parameters** | **Native *MsDps2*(small dataset)** | **MsDps2-DNA** | **$N_{15}$MsDps2-DNA** |
|---|---|---|---|
| **Microscope type** | Talos Arctica 200 kV cryo- EM | Talos Arctica 200 kV cryo- EM | Talos Arctica 200 kV cryo- EM |
| **Camera** | K2 direct electron detector (DED) | K2 direct electron detector (DED) | K2 direct electron detector (DED) |
| **Electron gun** | Field emission gun | Field emission gun | Field emission gun |
| **Voltage (HT)** | 200 kV | 200 kV | 200 kV |
| **Electron dose** | 40 $e^{-}/Å^2$ | 40 $e^{-}/Å^2$ | 40 $e^{-}/Å^2$ |
| **Electron dose per frame** | 2 $e^{-}/Å^2$ | 2 $e^{-}/Å^2$ | 2 $e^{-}/Å^2$ |
| **Total number of frames** | 20 | 20 | 20 |
| **Å/$\boldsymbol{pix}$** | 0.72 | 1.17 | 0.92 |
| **Defocus range** | -0.75 to -2.25 μm | -0.75 to -2.25 μm | -0.75 to -2.25 μm |
| **Plunge freeze instrument** | Vitrobot Mark IV | Vitrobot Mark IV | Vitrobot Mark IV |
| **Data collection mode** | Counting mode | Counting mode | Counting mode |
| **Symmetry imposed** | T | | T |

| | | T | |
|---|---|---|---|
| **Number of movie files** | 404 | 1091 | 838 |
| **Number of particles in model** | 98467 | 18168,171947 | 27741,36434 |
| **Map resolution (Å)** | 2.9 Å | 3.2 Å | 3.46 Å |
| **FSC threshold** | 0.143 | 0.143 | 0.143 |
| **Map sharpening B factor (Å$^2$)** | -163.1 | -188.9 | -201.5 |
| **MolProbity score** | 1.36 | 1.22 | 1.62 |
| **Poor rotamers (%)** | 1.45 | 0.00 | 0.00 |
| **Ramachandran plot** | | | |
| **Favoured (%)** | 99.42 | 98.79 | 99.13 |
| **Allowed (%)** | 0.58 | 1.21 | 0.87 |
| **Outliers (%)** | 0.00 | 0.00 | 0.00 |
| **EM Ringer score** | 3.43 | 2.28 | 1.82 |
